# Systematic optimization of Cas12a base editors in wheat and maize using the ITER platform

**DOI:** 10.1101/2022.05.11.491340

**Authors:** Christophe Gaillochet, Alexandra Pena Fernandez, Vera Goossens, Katelijn D’Halluin, Andrzej Drozdzecki, Myriam Shafie, Julie Van Duyse, Gert Van Isterdael, Camila Gonzalez, Mattias Vermeersch, Jonas De Saeger, Ward Develtere, Dominique Audenaert, David De Vleesschauwer, Frank Meulewaeter, Thomas B. Jacobs

## Abstract

The ever-increasing number of CRISPR components creates a significant burden when developing new genome engineering tools. Plant biotechnology in particular has few high-throughput options to perform iterative design-build-test-learn cycles when creating new gene-editing reagents. We have established ITER (Iterative Testing of Editing Reagents) based on arrayed protoplast transfections and high-content imaging, allowing one optimization cycle – from design to results– within three weeks. We validated ITER in wheat and maize protoplasts using Cas9 cytosine and adenine base editors. Given that previous LbCas12a-ABEs have low or no activity in plants, we used ITER to develop an optimized LbCas12a-ABE. We show that the sequential improvement of five components –NLS, crRNA, LbCas12a, adenine deaminase and linker– led to a remarkable increase in ABE activity from almost undetectable levels to 40% on an extrachromosomal GFP reporter. We confirmed the activity of LbCas12a-ABE at endogenous targets and in stable wheat transformants and leveraged these improvements to develop a highly mutagenic LbCas12a nuclease and LbCas12a-CBE. Our data show that ITER is a sensitive, versatile, and high-throughput platform that can be harnessed to accelerate the development of genome editing technologies in plants.

## Introduction

Base editors (BE) are incredibly useful biotechnological tools that generate precise nucleotide substitutions at specific DNA loci. There are currently two predominant types: cytidine (CBE) and adenine base editors (ABE). CBEs are a fusion of a cytidine deaminase domain to a catalytically inactive Cas nuclease or Cas nickase. A variety of cytidine deaminases have been used for base editing including APOBEC1 (A1), A3A, A3B, PmCDA1, AID, and their derivatives [1]. CBEs catalyze the deamination of cytidines into uracil on the non-target DNA strand, ultimately creating a C-to-T mutation [2]. CBEs also create C-to-G transversions and indels as byproducts, therefore many CBEs contain a uracil glycosylase inhibitor (UGI) to increase accuracy [2,3]. ABEs are derived from an evolved *Escherichia coli* adenine deaminase, TadA, that catalyze adenine into inosine which is repaired as guanine, leading to A-to-G transitions [4]. Unlike CBEs, the repair products are more accurate and fewer indels are observed [5].

CBEs and ABEs have been used in a wide range of species [6,7]. Cas9 has been the predominant nuclease, which has limited the target range since the protospacer-adjacent motif (PAM; 5’-NGG-3’) and deamination window control the target space. Numerous groups have utilized Cas9 PAM variants to overcome this limitation [8–12]. In contrast, few reports have utilized Cas12a for base editors with low to no activity [13–18] and only proof-of concept Cas12a-BEs have been developed for plants [17,19–21]. Therefore, it is highly desirable to generate an efficient Cas12a-BE to increase the targeting space (5’-TTTV-3’ PAM). Such a reagent would also be an attractive alternative to Cas9 due to a simpler intellectual property landscape [22].

The optimization of genome editing tools such as base editors requires testing large numbers of configurations (architectures) and targets [2,4,23]. To facilitate the systematic development of BEs specifically for plants, we developed ITER (**I**terative **T**esting of **E**diting **R**eagents), a high-throughput, multi-well plate-based system for fast, versatile, and quantitative results to iteratively test and optimize novel genome editing technologies. We first validated ITER by testing a variety of Cas9-CBEs and ABEs in wheat and maize protoplasts. We then systematically developed and optimized an LbCas12a-ABE by modifying various NLS, linker, and crRNA components in five iteration cycles. We quickly go from near-background levels (0.5%) of ABE activity, to as high as ~40% for an extrachromosomal GFP reporter and 10-20% for endogenous targets in wheat and maize. We confirmed these novel ABEs in stably-transformed wheat lines where the most active generated edits in ~30-50% of independent events. These optimizations are generally applicable as they also increase indel rates from ~10% to ~80% with nuclease-active LbCas12a and allow for LbCas12a-CBE activity. Thus, we demonstrate that ITER is an effective, high-throughput platform to accelerate the development of genome editing tools in plants and present highly optimized architectures for LbCas12a.

## Results

### ITER platform establishment and validation

We aimed to develop a platform that combines arrayed protoplast transfections with high-content imaging to efficiently test and optimize a wide variety of genome editing tools (Fig. 1a). We first scaled up the number of wheat protoplast transfections by reducing the number of cells and volume per transfection (5 × 10^5^ cells in 500 μl to 1 × 10^5^ cells in 100 μl) and then arraying the transfections in a 96-well plate format for streamlined analysis using the OPERA® High-Content Screening System for automated cell detection and fluorescence analysis (Methods, Fig. 1a and Additional file 1: Fig. S1a). After cell segmentation and quantification based on two fluorescent reporters, p35S:mCherry-NLS and pZmUBI:GFP-NLS, on average 1,000 to 1,500 transfected cells were detected per sample with a transfection efficiency of 40-65% (Additional file 1: Fig. S1a and S1c). Importantly, co-transfection efficiency is effectively 100% and homogeneously distributed throughout the plate (Fig. 1b). We then adapted the workflow for etiolated maize protoplasts by segmenting cells only on nuclei expressing mCherry, demonstrating that ITER can also be used in multiple plant species to process cells with or without chlorophyll (Methods and Additional file 1: Fig. S1d, e).

**Fig. 1:**
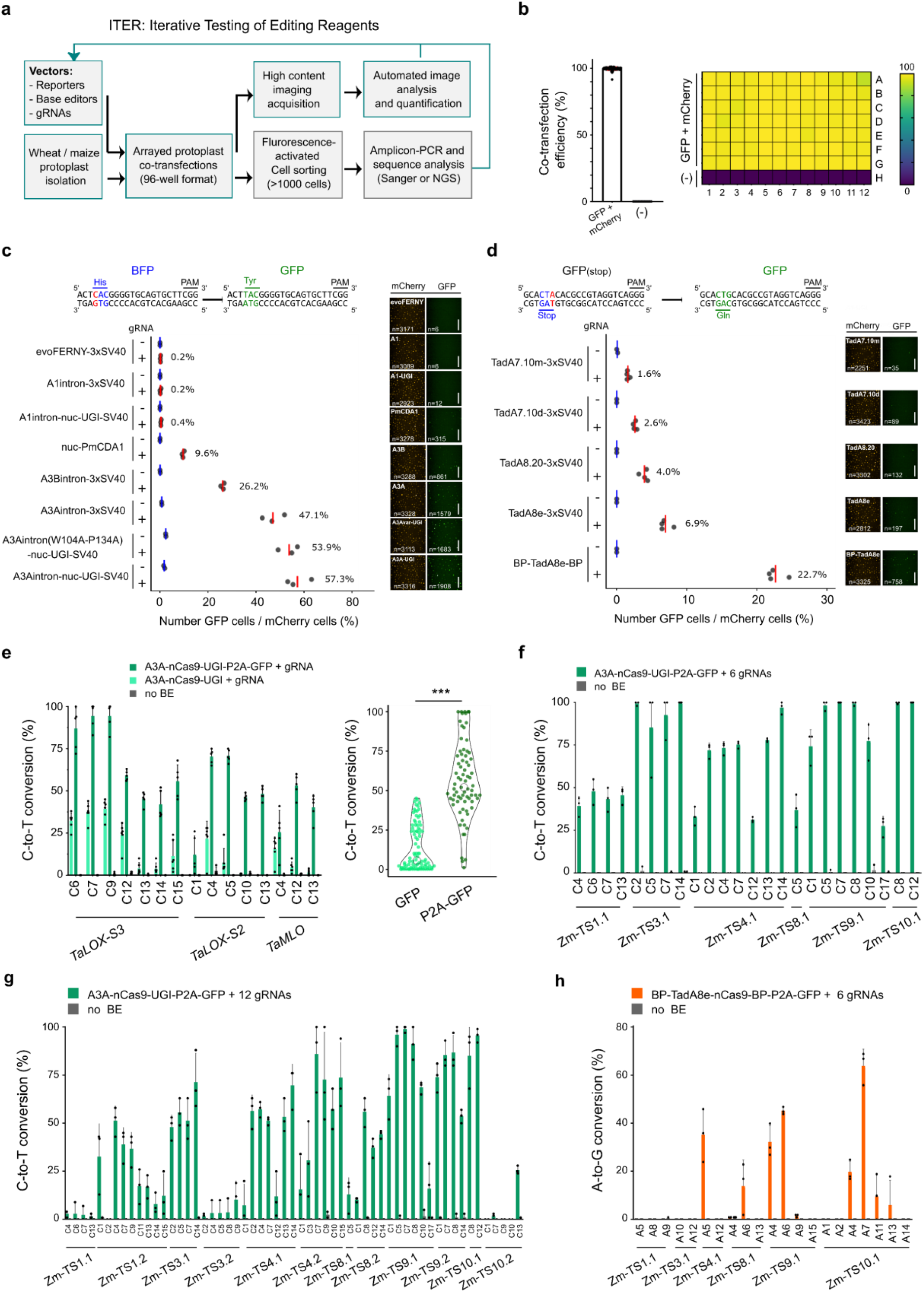
ITER is an efficient platform for testing base editing in wheat and maize protoplasts. **a,** Schematic illustration of ITER. Wheat or maize protoplasts are co-transfected in 96-well format. Cells are imaged and analyzed by automated high content imaging. Resulting data are interpreted and inform the subsequent optimization cycle. Validation of base editor activity at endogenous targets is performed by genotyping after Fluorescence-activated cell sorting (FACS). **b,** Co-transfection efficiency as measured by the number of GFP over the number of mCherry positive cells in wheat protoplasts. Negative controls (−) do not contain vector DNA. The grid shows the 96-well plate positions and the gradient colors coding represent co-transfection efficiencies. **c-d,** Base editing efficiencies for multiple nCas9-CBE (c) or nCas9-ABE (d) configurations in wheat protoplasts. Efficiencies were measured as the percentage of cells showing BFP to GFP conversion for C-to-T (c) or GFP recovery for A-to-G (d). Red and blue lines represent mean efficiency of cells transfected with or without gRNA, respectively. Representative pictures show mCherry or GFP fluorescence in protoplasts. n indicates the total number of cells measured after image segmentation in the mCherry or GFP channel. Scale bars: 200 μm. **e,** Multiplex C-to-T base editing of A3A-nCas9-UGI or A3A-nCas9-UGI-P2A-GFP at three wheat targets. One sub-genome was analyzed by Sanger sequencing and EditR analysis. Asterisks represents significant difference in efficiency as measured by Student’s-t test (***: p<0.001). **f-g,** Multiplex C-to-T base editing of A3A-nCas9-UGI at 6 (f) or 12 (g) maize targets. **h,** Multiplex A-to-G base editing efficiency of BP-TadA8e-nCas9-BP ABE at six maize targets. Transfection of the cells with a GFP expressing vector (no BE) was used as negative control in e-h. X-axis in e-h indicates targeted Cytosines (e-f) or Adenine (h) at different positions along the protospacer, with the PAM-distal base being position 1. Editing rates were calculated from three to six independent biological replicates that are depicted as dots on plots (c-h).

We tested ITER by comparing the activity of eight Cas9-CBEs with different components; the cytidine deaminase, NLS variants and with or without UGI (Fig. 1c and Additional file 1: Fig. S2a). We co-transfected three vectors: 1) a BFP-to-GFP reporter, 2) a CBE and 3) a gRNA targeting BFP [24,25]. As a negative control, the reporter and the CBE were co-transfected to monitor non-specific base editing. Editing efficiencies (the ratio of GFP to mCherry positive cells) ranged from 0.2% (n=6/3171) to 57.3% (n= 1908/3316), depending on the architecture, showing that we can detect C-to-T editing with high sensitivity (Fig. 1c). Interestingly, editing efficiencies correlated well with average GFP intensities, indicating that the number and intensity of GFP positive cells can also inform on BE activity (Fig. 1c and Additional file 1: Fig. S2b). Overall, the relative activities of the different deaminases without UGI matched those reported in the literature with A3A>A3B>PmCDA1>A1 and A3A with a C-terminal UGI leading to the highest editing rates [23,26,27], confirming that our platform can be used to reliably compare CBE activities. We next tested whether we could monitor ABE activity using a GFP(stop) to GFP reporter where a CAG codon is replaced by a TAG stop codon [28] and compared the activity of five nCas9-ABEs that carry different adenine deaminases and NLS variants. Editing efficiencies ranged from 1.6%-22.7%, with the Tad8e variant containing N- and C-terminal BP NLS being the most active (Fig. 1d).

We transfected a subset of the nCas9-CBEs and ABEs into maize protoplasts to test ITER in a different crop species. We reliably detected BE activity of both reporters and observed efficiencies were higher than in wheat, but showed the same overall trends (Fig. 1c, d and Additional file 1: Fig. S2d, e). We also confirmed the activity of A3A-PBE and ABE-7 and observed that ABE-7 displayed higher efficiency than a nearly identical ABE built in our laboratory (Fig. S1d; Additional file 1: Fig. S2f), showing that our platform can be used to quickly validate and compare reagents from other laboratories [27,28] (Additional file 1: Fig. S2d, f). Together, these results show that ITER is a rapid, sensitive and versatile platform for assessing BE activity in wheat and maize protoplasts.

Testing BE activity with a single target site in a reporter plasmid may bias results for specific target sequence contexts. Therefore, we also added a module to ITER to compare the activity of multiple BEs at endogenous targets. We first scaled up Fluorescence-Activated Cell Sorting (FACS) and genotyping strategies to increase the number of sorted wheat samples from 4-6 samples to 15-25 per hour (Methods and Additional file 1: Fig. S3a, b). We selected the CBE A3A-nCas9-UGI, which performed best in our GFP reporter assays, and measured editing at the *TaLOX2-S3* target with Sanger sequencing after sorting 1,000 or 2,000 cells on GFP (Additional file 1: Fig. S3c). Base editing ranged from 10 to 50% regardless of the number of sorted cells, indicating that we can reliably measure BE activity at endogenous targets using as few as 1,000 sorted cells (Additional file 1: Fig. S3c). We did not detect C-to-G nor C-to-A editing, indicating that A3A-nCas9-UGI predominantly introduces C-to-T mutations in protoplasts.

We next tested if enriching for BE-expressing cells by utilizing a ribosomal skipping P2A reporter could increase sensitivity (Fig. 1e). We co-transfected base editors with and without P2A-GFP-NLS, with three gRNAs in multiplex targeting *TaLOX-S3, TaLOX-S2* and *TaMLO* and sorted 2,000 cells on GFP. Analysis of the three targets detected a significant increase in editing activity for the A3A-nCas9-UGI-P2A-GFP-NLS relative to the A3A-nCas9-UGI (from 13.6% to 56.3%; Mann-Whitney test: p<0.001) and a broader editing window (from C4-C12 to C1-C15) (Fig. 1e). As these results confirmed that the P2A-GFP-NLS module can increase sensitivity, we used the P2A-GFP-NLS module in our subsequent BE architectures. We then increased the number of gRNAs to six and 12 in maize cells using gRNA arrays (Methods). For the 6-gRNA arrays, we detected C-to-T conversions at all targets with values ranging from 25-100% (Fig. 1f and Additional file 1: Fig. S3d) whereas for the 12-gRNA (both arrays co-transfected), we observed base editing at nine out of 12 targets and a 5%-100% decrease in efficiency compared to the 6-gRNA arrays (Additional file 1: Fig. S3e).

To confirm ABE activity at endogenous targets, we selected BP-TadA8e-nCas9-BP, which showed the highest A-to-G conversion in our GFP reporter assay. Only one out of four sites was edited in simplex for wheat (Additional file 1: Fig. S3f). For maize, four out of six targets were edited for array1 (Fig. 1h) and three out of six for array2, with editing efficiencies ranging from 5 to 65% (Additional file 1: Fig.S3g). When combining the two arrays, we detected five out of twelve targets edited at 5 to 40% efficiency and editing windows ranged from A4 to A10 (Fig. 1h and Additional file 1: Fig. S3f, h). These results demonstrate that we can efficiently detect multiplex C-to-T or A-to-G base editing but also suggest that increasing the number of gRNAs negatively correlates with BE activity at individual targets in protoplasts.

### Development and optimization of Cas12a-ABE with ITER

We wanted to leverage ITER to develop a new class of base editors for plants. While Cas12a-BEs have been used in animal systems [13–16], these tools have not been widely adapted to plants [17,19,21]. This stands in contrast to Cas9-BEs and nuclease-active Cas12a which are widely used in both fields [29,30]. We started by building fluorescent reporters to detect LbCas12a-CBE or ABE activity. For the CBE reporter, we used BFP with a T63S substitution to add a Cas12a PAM (5’-TTTC-3’; Methods, Additional file 1: Fig. S4a). For the ABE reporter, the R110 (CGA) GFP codon was mutated into a stop codon (*; TGA) (Methods, Additional file 1: Fig. S4b). We first tested the activity of A3A-dCas12a-UGI and four versions of dCas12a-ABE, including TadA8e-32aa-dCas12a-3xSV40, using a wheat codon-optimized LbCas12a (Additional file 1: Fig. S4a, b). Editing levels for CBEs and ABEs were similar to negative controls without crRNAs, indicating that the measured activity was non-specific (Additional file 1: Fig. S4a, b), possibly due to unspecific deaminase activity or background fluorescence of the reporter [31]. We decided to focus on the dCas12a-ABE system as we were limited to only one reporter to test CBE activity. Using a new GFP reporter (Q70*/R74K) [19], we repeatedly detected a low but consistent signal ranging from 0.6% to 0.8% (n>8000) for TadA8e-32aa-dCas12a-3xSV40 whereas no signal was detected for the negative control nor the other TadA variants (Fig. 2a,c and d1 and Additional file 1: Fig. S4c). These results provided a starting point for optimization with TadA8e-32aa-dCas12a.

**Fig. 2:**
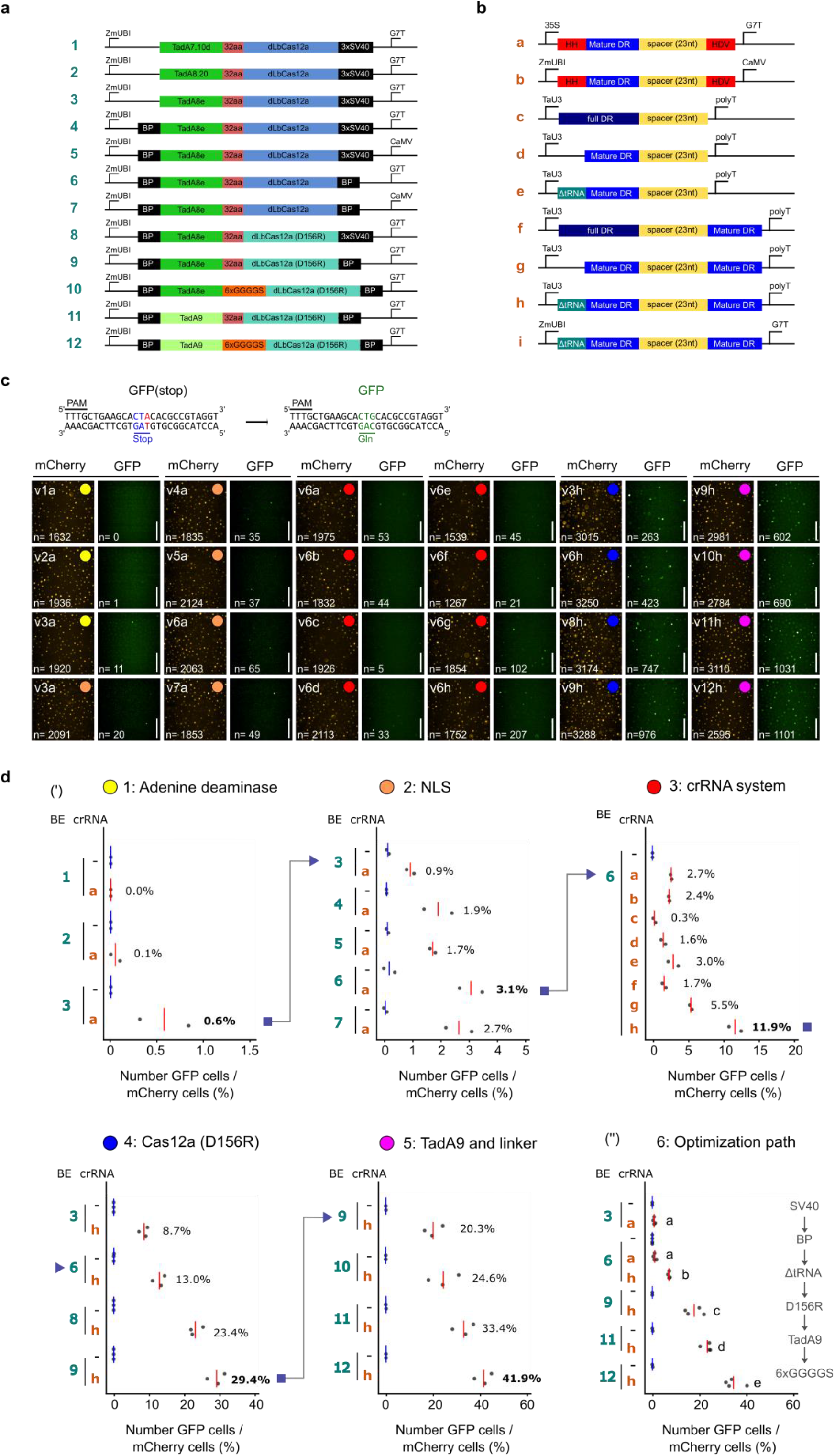
Stepwise optimization of Cas12a-ABE using ITER. **a-b,** Schematic illustration of 12 LbCas12a-ABEs (a) and nine crRNA (b) architectures tested in this study. **c,** Representative pictures showing mCherry and GFP positive cells measured during the optimization campaign of LbCas12a-ABE in wheat protoplasts. A-to-G base editing of the GFP reporter recovers GFP fluorescence. Colored dots represent different iterations and match measurements in d. Scale bars: 200 μm. **d**, Editing efficiencies of Lb-Cas12a ABE as measured by the rate of GFP recovery in wheat cells. BE and crRNA architectures are referred to as numbers and letters, respectively, as shown in a-b. Red and blue lines represent mean efficiency of cells transfected with or without crRNA, respectively. (‘) 5 ITER cycles were conducted by sequentially testing adenine deaminases, NLSs, crRNA systems, Cas12a variants and protein linkers. Best candidates obtained at individual iterations were used as baseline in the following iteration and is depicted with a blue arrow head. (“) Key Cas12a-BE architectures obtained along the optimization path were compared side by side. Components leading to increased activity are shown at the right of the panel. Editing rates in d were calculated from 2-4 independent biological replicates that are depicted as dots on plots. Statistical significance is calculated with one-way ANOVA test followed by Tukey HSD post-hoc test at threshold p<0.05.

We next tested two NLSs and their position on the ABE together with the G7 terminator or CaMV terminator as this terminator is used on the more active PABE-7 [28] (Fig. 2a, c, d2). The BP NLS on both the N- and C-terminus increased base editing to 3.1%, suggesting that improving nuclear localization of the complex increases activity as previously shown with our Cas9 results (Fig. 1c) and other genome engineering tools [32]. We then compared eight different crRNA architectures with either the standard Pol II-driven mature direct repeat (DR) flanked by ribozymes, Pol III-driven full DR, mature DR, or truncated tRNA [33] (Fig. 2b). Strikingly, the TaU3:tRNA-DR-spacer-DR architecture (h) led to a 4-fold increase in efficiency (2.7 to 11.9%) compared to the standard p35S:HH-spacer-HDV (Fig. 2c, d3).

In the fourth iteration, we tested the LbCas12a D156R mutation as this variant has high indel activity in *Arabidopsis* [34]. This variant increased activity to 29.4% GFP-positive cells (Fig. 2c, d4). In the fifth iteration, we tested a more active TadA, TadA9 [35], and a 6xGGGGS linker as this was reported to increase dCas12a-CBE activity in maize [21]. Both variants further increased the activity of the Cas12a-ABE to 42% efficiency (Fig. 2c, d5). Finally, we tested whether we could further improve the crRNA system by using a Pol-II promoter driving the crRNA as previously reported [33] (Fig. 2b, c). We observed lower editing efficiencies for ZmUBI:tRNA-DR-spacer-DR than for the TaU3 promoter (h) (Additional file 1: Fig. S4d), confirming that TaU3:tRNA-DR-spacer-DR was the best crRNA architecture from our panel.

We compared the successive BE architectures in a single ITER run to confirm the role of individual modifications on LbCas12a-ABE activity (Fig. 2c, d6 and Additional file 1: Fig. S4e). In line with our previous experiments, introducing the truncated tRNA DR-DR crRNA system, the LbCas12a(D156R) variant, TadA9 and the 6xGGGGS linker all led to significant increases in base editing efficiencies (One-way ANOVA, Tukey HSD: P<0.05). We also observed comparable stepwise increases in Cas12a-ABE activity in maize protoplasts (0.8 to 67.9%) indicating that the optimizations are transferable to other plant species (Additional file 1: Fig. S4f).

### Comparative analysis of Cas12a ABEs at endogenous sites and in transformed plants

After developing LbCas12a-ABEs with a reporter, we wanted to validate the key optimizations at endogenous targets. We first randomly selected six targets and confirmed v9h activity at five out of six targets in protoplasts. Simplex and multiplex editing levels ranged from 0.5-4% (Additional file 1: Fig. S5a, b). We then selected six key LbCas12a-ABE/crRNA architectures and assessed their activity at the four best targets in simplex (Fig. 3a). For the three best versions (v9h, v11h, and v12h) we detected A-to-G conversion at all four targets ranging from 0.5-9% with an editing window spanning A8 to A11. We also observed low indel frequencies (0.5% and 5%) at two targets with v9h (Additional file 1: Fig. S5e). Combining data points for all four targets revealed that introducing the truncated-tRNA DR-DR, LbCas12a (D156R), TadA9, and 6xGGGGS linker led to stepwise increases in editing efficiency (Kruskal-Wallis test; Dunn post-hoc test a P<0.05). We observed similar results when analyzing multiplex editing at the same four targets (Fig. 3b and Additional file 1: Fig. S5b).

**Fig. 3:**
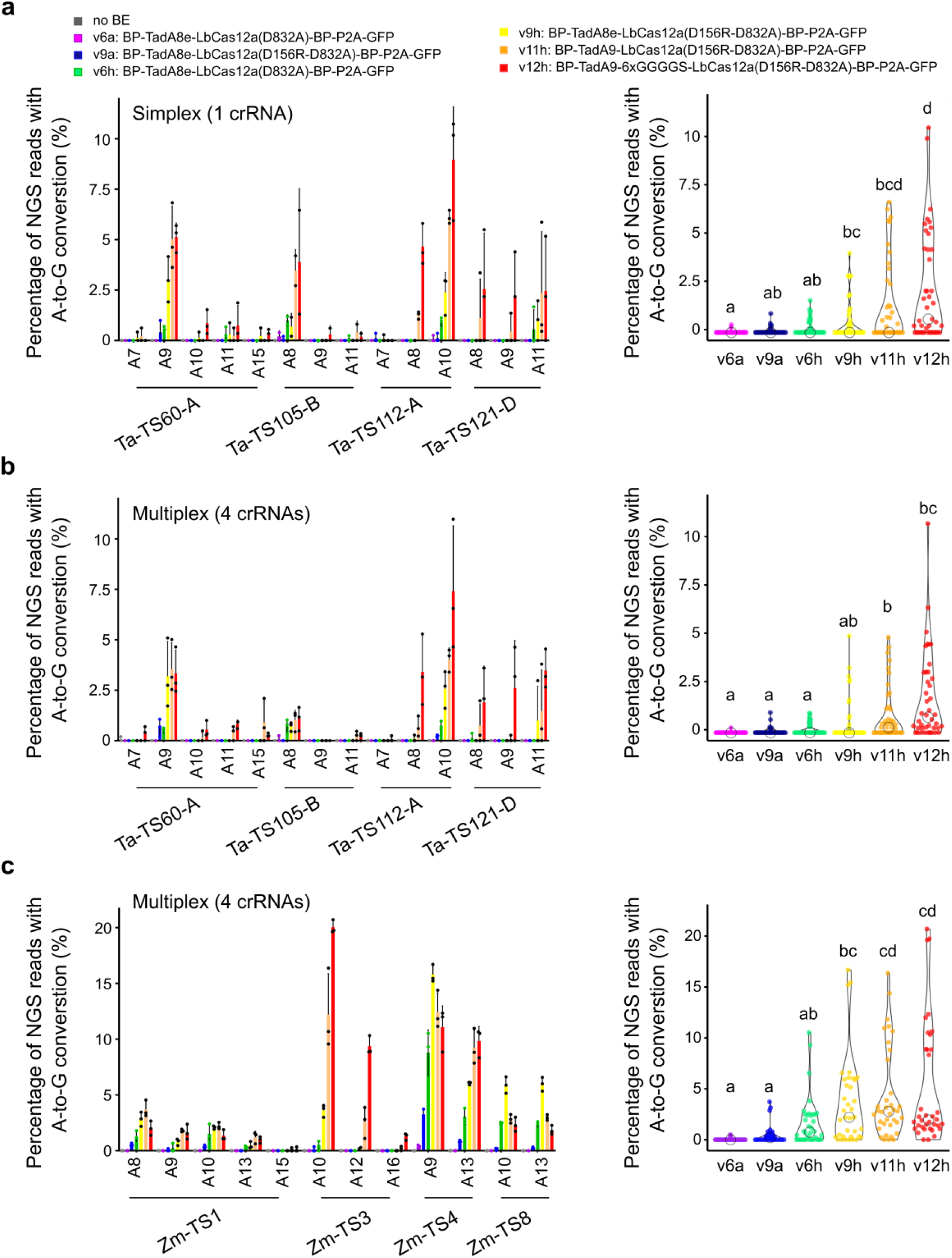
Validation of Cas12a-ABE activity at endogenous targets in wheat and maize. **a-c,** Simplex (a) or multiplex (b-c) base editing efficiencies of six Cas12a-ABE configurations as measured by the proportion of NGS reads converted from A-to-G in wheat (a-b) or maize (c). Different Cas12a-ABE versions are labeled with number and letters referring to Fig. 2a-b. Barplot displays A-to-G conversion rates at individual on-target target. x-axis indicates targeted adenine at different positions along the protospacer, with the PAM-adjacent base being position 1. Editing rates were calculated from 2 or 3 independent biological replicates that are depicted as dots on barplots. Violin plot represent pooled efficiencies at all targets for individual base editor versions. Statistical significance is calculated by Kruskal-Wallis test followed by Dunn post-hoc test at threshold p<0.05.

To check off-target activity, we leveraged the allohexaploid nature of the wheat genome and selected targets where one subgenome perfectly matched the crRNA sequence and the two homeologs contained mismatches (Methods, Fig. 3a, b; Additional file 1: Fig. S5c, d). Mismatches in the seed region of the spacer strongly decreased ABE activity, whereas mismatches at the PAM-distal end led to editing efficiencies ranging from 2.5-7% (Additional file 1: Fig. S5c, d).

We then analyzed the same six LbCas12a-ABE and crRNAs architectures in maize at four target sites in multiplex. Consistent with the results in wheat, LbCas12a-ABE v9h, v11h, and v12h performed best with editing efficiencies ranging from 0.5-20% (Fig. 3c; Additional file 1: Fig. S5f; Kruskal-Wallis test; Dunn post-hoc test a P<0.05), further validating the BE optimization campaign in protoplasts.

We tested the v9h and v11h architectures be generating transformed plants with the two best crRNAs via co-bombardment (TS60 and TS112; Fig. 4). We regenerated 154 and 184 independent T0 plants for v9h and v11h, respectively, and evaluated base editing with NGS. The percentage of plants carrying A-to-G edits at primary positions (A7 and A9 for TS60-A or A8 and A10 for TS112-A) ranged from 2.6-34% for v9h and 11.5-46.7% for v11h (Fig. 4a, b). Consistent with the protoplast results, v11h was significantly more active than v9h (1.4 to 4.7 fold increase, p<0.05 z-score test for two proportions; Fig. 4a, b). The majority of the mutated lines were heterozygous with 13 and 29% of the lines containing heterozygous on-target edits for v11h at TS60-A and TS112-A, respectively. Homozygous lines were also recovered at 3–18% for the primary positions (Fig. 4a, b and Additional file 1: Fig. S6b). In addition, base editing was observed at secondary positions (A10, A11 and A15 for TS60-A and A6, A7 for TS112-A) and no indels were detected (Fig. 4a, b). Altogether, v9h and v11h activity at TS60-A and TS112-A created a panel of 18 and 37 unique genotypes, respectively (Fig. 4c).

**Fig. 4:**
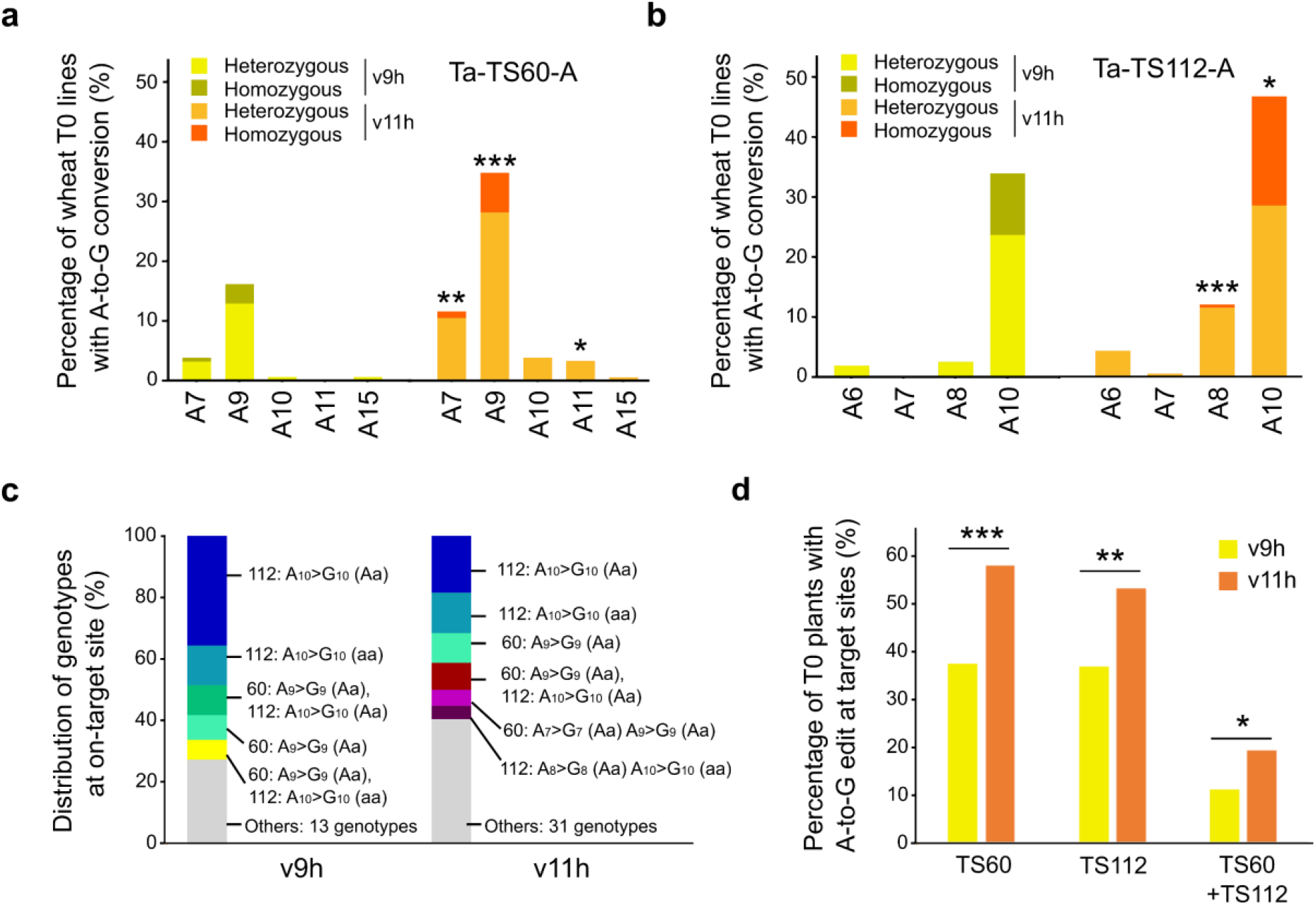
Validation of Cas12a-ABE activity in stable wheat plants. **a-b**, Percentage of T0 wheat plants with A-to-G base editing at independent positions of the target as measured by NGS (v9h: n=154; v11h: n=184). Two Cas12a-ABEs (v9h and v11h) are compared at TS60-A (a) and TS112-A (b). Percentage of T0 wheat plants with heterozygous (editing rates between 25% and 75%) and homozygous mutations (higher than 75%) are displayed. **c**, Frequency of genotypes generated by Cas12a-ABE v9h and v11h. A>G indicates base editing measured at the target position. Aa and aa denote heterozygous and homozygous base editing on targets, respectively. **d**, Percentage of T0 wheat plants carrying at least one mutation in at least one subgenome at TS60 or TS112 loci. Asterisks depict significant difference in editing efficiency between v9h and v11h as measured by z-score test for two proportions (*: p<0.05; **: p<0.01; ***: p<0.001).

We analyzed off-target edits and detected a significant increase in A-to-G conversion for v11h compared to v9h, consistent with previous results (Additional file 1: Fig. S6a; p<0.01; z-score test for two proportions). Around 30% of the plants transformed with v9h or 50% with v11h carried at least one base edit in at least one of the off-targets at TS60 or TS112 (Fig. 4d and Additional file 1: Fig. S6b). Furthermore, double mutants for TS60 and TS112 were generated at a rate of 11% for v9h and 19% for v11h (Fig. 4d). Together, these results show that both Cas12a-ABEs can be used to efficiently introduce multiplex base edits in stable wheat lines without indels.

### Transfer of improved components to the LbCas12a toolset

We wondered whether the key optimizations of LbCas12a-ABEs could more broadly improve the LbCas12a toolset. We built three CBEs with A3A based on architectures v3, v11 and v12 and co-transfected wheat protoplasts with either the 35S:HH-spacer-HDV (a) or TaU3:tRNA-DR-spacer-DR (h) crRNA systems (Fig. 2a, b). We measured multiplex base editing at four endogenous targets and detected C-to-T substitutions at all sites with editing efficiencies ranging from 1-23% and an editing widow spanning C6-C17 (Fig. 5a). In line with our ABE results, CBEs architectures corresponding to v11h and v12h performed best (Kruskal-Wallis test; Dunn post-hoc test a P<0.05) (Fig. 5a). Interestingly, we also observed high editing rates at two off-targets (TaTS13-A and TaTS60-D) despite the presence of mismatches in the 3’ region of the crRNA (Additional file 1: Fig. S7a).

**Fig. 5:**
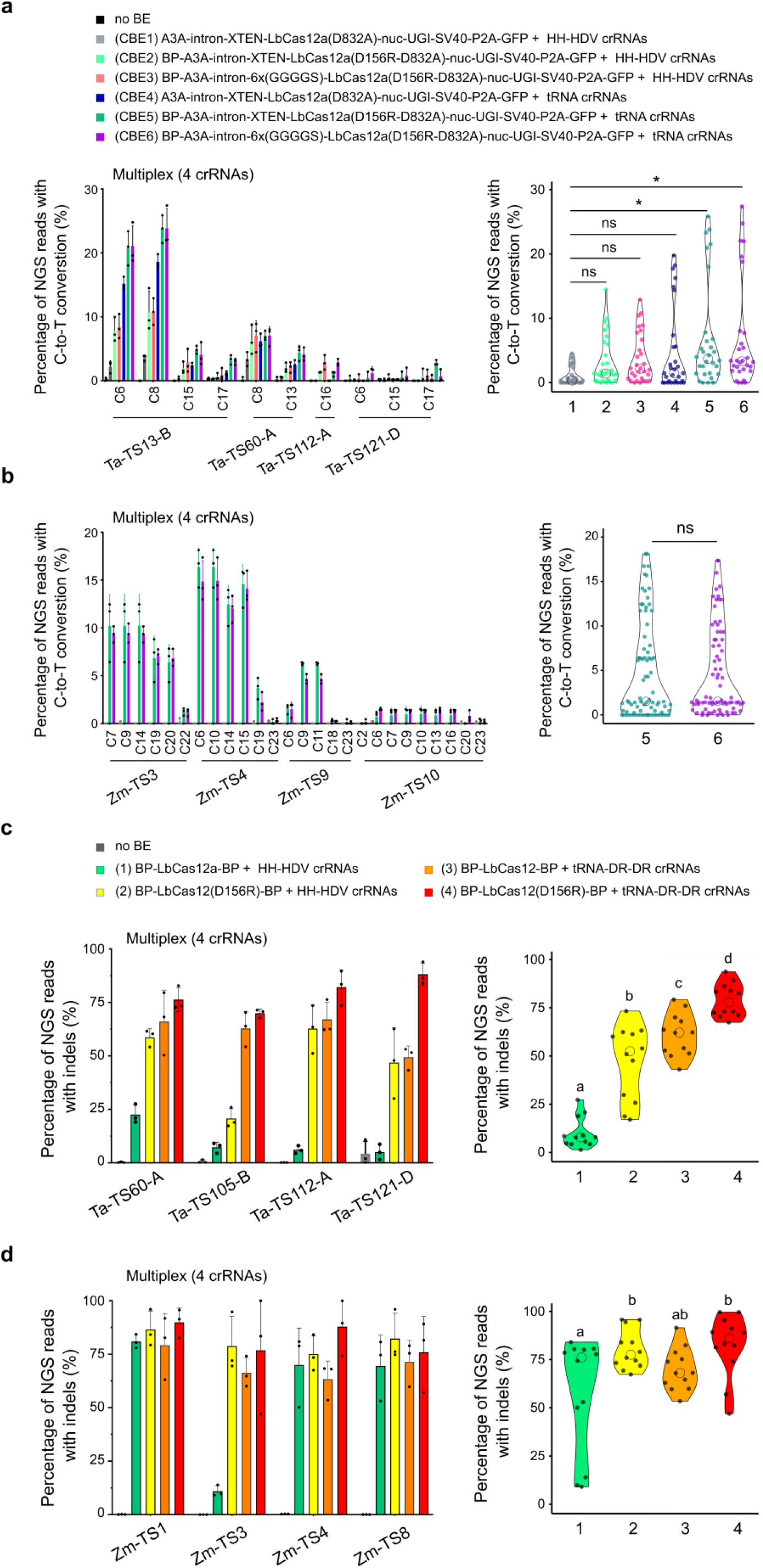
Building a Cas12a toolset with optimized components. **a-b,** Multiplex C-to-T base editing efficiencies of six LbCas12a-CBE versions in wheat (a) or two versions in maize (b). The x-axis indicates targeted cytosines at different positions along the protospacer, with the PAM-adjacent base being position 1. Violin plots represent pooled efficiencies at all targets for individual base editor architectures. **c-d**, Indel rate for four versions of LbCas12a nuclease in wheat (c) or maize (d). Barplots display indel rates at individual homeolog on-targets. Violin plots represent pooled indel rates at all targets for individual LbCas12a architectures. Editing rates were calculated from three independent biological replicates that are depicted as dots on barplots (a-d). Statistical significance in a is calculated by Kruskal-Wallis test followed by Dunn post-hoc test (*: p<0.05); in b using Mann-Whitney test; in **c-d** by two-way ANOVA test with Tukey HSD post-hoc test at threshold p<0.05.

We confirmed the activity of the two best CBEs (CBE5 and CBE6) in maize protoplasts by measuring multiplex base editing at four targets and detected C-to-T substitution at all sites ranging from 1-15% (Fig. 5b). The editing window spanned C6-C22, consistent with previous report for A3A [27]. We did not observe a significant difference in activity between CBE5 and CBE6 architectures suggesting the linkers do not change CBE activity in maize protoplasts (Mann-Whitney test, P<0.05; Fig. 5b).

We also compared indel rates for nuclease-active LbCas12a and LbCas12a(D156R) with either the 35S:HH-spacer-HDV (a) or TaU3:tRNA-DR-spacer-DR (h) at four wheat targets in multiplex (Fig. 4c). Consistent with the BE results and previous reports in plants and animals [33,34,36], both LbCas12a(D156R) and truncated tRNA-DR-DR increased LbCas12a mutagenic activity from 10% to 47% and 65%, respectively. Combining both components significantly increased indel formation to 79% without changing the indel pattern (Two-way ANOVA, Tukey HSD: P<0.05; Fig. 5c and Additional file 1: Fig. S7b). Off-target analysis additionally confirmed that mismatches in the seed region of the Cas12a spacer dramatically decrease LbCas12a mutagenic activity whereas mismatches at the 3’ end do not (Additional file 1: Fig. S7b). In line with our observations in wheat, indel rates at four targets in maize for these four LbCas12a architectures showed that the best architecture also performed well with an average indel rate of 82% (Fig. 5d). The three other architectures also induced high indel rates (57-80%), suggesting that these optimizations are less crucial in maize (Fig. 5d). Altogether, we show that key optimization components gained from our LbCas12a-ABE optimization campaign could be broadly applied to develop efficient LbCas12a-CBE and LbCas12a nuclease systems.

## Discussion

We have established ITER, a sensitive, versatile and high-throughput method to test and quantify gene editing outcomes using a fully automated imaging and analysis pipeline. An iterative cycle can be performed in approximately three weeks where new entry clones are created in week one, the panel of vectors are assembled and validated in week two and plasmid scale up, transfection and analysis occur in week three. ITER automatically normalizes for transfection efficiency and can efficiently process multiple replicates to confidently detect low-efficiency signals even when the transfection efficiency variance between samples can be high (~2-fold). As a result, ITER is able to reliably detect as few as 3-14 GFP positive cells with an average threshold efficiency of 0.6% (8 replicates: Fig. 2d and Additional file 1: Fig. S4c).

The use of fluorescent reporters does have limitations; high background noise was observed in one of the GFP reporters, possibly due to gRNA-independent DNA and/or RNA editing [31,37,38]. Furthermore, measuring editing at a single target site limits the types of repair outcomes that can be evaluated (e.g. editing window, bystander edits and sequence context). As a complementary approach, we targeted endogenous genes and optimized FACS methods to select cells highly expressing the reagents, thereby increasing sensitivity. A relatively low number of cells (~1,000) can be used to detect mutations, reducing sorting time which becomes a limiting factor when scaling up.

We used ITER to conduct an Cas12a-ABE optimization campaign and identified key components leading to improved ABE, CBE and nuclease activity in both wheat and maize. Notably, only a handful of reports have only provided proof-of-concept demonstrations of Cas12a-BE vectors in plants [17,19–21]. We think that this is likely due to the low activity in plant cells with standard Cas12a designs. The results from the start of our optimization campaign support this as we were unable to detect any base editing with A3A-dLbCas12a-UGI and only low efficiencies for TadA8e-dLbCas12a. The use of ABE variants increased activity, however, this did not occur at all targets indicating that the target sequence context and/or crRNA might also impact editing efficiency. Our results using alternative crRNA architectures largely agree with those of Zhang et al., 2018 [33]; the inclusion of the mature DR on both ends of the crRNA and the 5’ truncated tRNA motif increased LbCas12a activity. However, in contrast to the previous report, Pol III-driven crRNAs were more effective than Pol II crRNAs.

Editing levels of our LbCas12a-ABE are notably lower than those with nCas9. We think this is likely due to the nickase activity of Cas9 which promotes the use of the non-target strand as a template in DNA mismatch-mediated repair. There is currently no comparable Cas12a nickase as Cas12a sequentially cuts the non-target and then the target strand [39] and nickase mutants only cut the non-target strand. Therefore, we and others [16] hypothesize that creating a Cas12a variant that can nick only the target strand would increase the efficiency of Cas12a-ABEs and -CBEs.

## Conclusions

Here, we have demonstrated the effectiveness of ITER to systematically optimize base editors in two important crops, wheat and maize. Starting from nearly no activity, the combined stepwise optimizations allowed us to create a highly effective LbCas12a-ABE. Together with the LbCas12a-CBE, this toolset expands the target range of base editing to T-rich genomic regions in plants. As protoplasts are readily obtainable from many plant species, we also expect that ITER will be broadly applicable across plants and envision that it will be used to rapidly optimize a variety of genome editing tools in plants such as HDR [40], prime editing [41,42] or even new types of base editing [43,44].

## Materials and methods

### Cloning

PCR was performed using Q5® High-Fidelity DNA Polymerase (M0491, NEB) with DNA oligos from Integrated DNA Technologies (IDT). PCR products were gel purified using Gel Purification Kit (D4002, Zymo Research). To generate entry vectors, DNA fragments were inserted into BsaI-digested GreenGate empty entry vectors [45] via Gibson assembly (2x NEBuilder Hifi DNA Assembly Mix, NEB) or restriction ligation with T4 DNA ligase (NEB). Base editors, gRNAs and fluorescent reporter vectors were assembled using Golden Gate cloning (30 cycles (37°C, 5 min; 16°C, 5 min); 50°C for 5 min; 80°C for 5 min) with BsaI or BbsI [45]. Vectors were transformed by heat-shock transformation into DH5α *E.coli* or One Shot™ ccdB Survival™ competent cells (Thermo Fisher Scientific). Cells were plated on lysogeny broth medium containing 100 μg mL^−1^ carbenicillin, 100 μg mL^−1^ spectinomycin, 25 μg mL^−1^ kanamycin or 40 μg mL^−1^ gentamycin, depending on the selectable marker. Plasmids were isolated (GeneJET Plasmid Miniprep kit, Thermo Fisher Scientific) and confirmed by restriction enzyme digestion and/or Sanger sequencing (Eurofins, Mix2seq). All plasmids are described in Additional file 6. All primers are listed in Additional file 7.

#### Entry clones

The cytidine deaminase sequences of PmCDA1, APOBEC3A and APOBEC3B were synthesized on the BioXP3200 DNA synthesis platform (Codex DNA) based on published sequences [24,46] and inserted into empty entry vectors by Gibson assembly. APOBEC1 and UGI entry clones were previously described [47]. evoFERNY was PCR amplified from the Addgene plasmid #125615 [48] and inserted into empty entry vectors by Gibson assembly.

The (GGS)_5_ linker on the APOBEC1 entry clone was replaced with the XTEN linker by Gibson assembly. Due to apparent toxicity of APOBEC vectors in *E.coli*, the *Solanum tuberosum* IV2 intron [49] was added to APOBEC1, A3A and A3B via Gibson assembly. The intron insertion site was designed using NetPlantGene [50].

TadA7.10d was synthesized on the BioXP3200 DNA synthesis platform (Codex DNA) based on the published sequence [4], TadA8e (Addgene #138489)^14^ and Tad8.20m (Addgene #136300)^50^ were PCR amplified and inserted into empty entry vectors by Gibson assembly.

The LbCas12a(D832A) sequence was codon-optimized for wheat using a BASF proprietary software tool and synthesized (Twist Biosciences). Three synthesized fragments were cloned into an entry vector using Gibson assembly. The LbCas12a(D156R-D832A) variant was generated by site-directed mutagenesis PCR via Gibson assembly. The 3xSV40-NLS and BP-NLS were based on published sequences [14,28] and cloned by annealed oligos followed by ligation. Nuc-UGI-SV40 was amplified from A3A-PBE (Addgene #119768) [27] and cloned into an entry vector via Gibson assembly. The CaMV terminator was isolated from the PABE-7 plasmid (Addgene #115628) [28] and cloned into an entry vector via Gibson assembly.

#### gRNA and crRNA

*TaLOX-S2* and *TaLOX-S3* and *TaMLO* Cas9 targets were previously reported [27,52]. Maize Cas9 and Cas12a targets were designed on genes controlling yield [53]. Cas9 gRNAs were cloned via Golden Gate with BbsI using annealed oligos into the entry vectors (p01356-p01361) as previously described [54]. The 6-gRNAs arrays were assembled using Golden Gate with BsaI of the individual TaU3:gRNA entry clones. Wheat Cas12a targets were selected using an in-house script on the Fielder genome (BASF) so that the crRNAs would perfectly match one subgenome and contain mismatches with the other two. Cas12a crRNAs vectors were cloned by amplifying a TaU3 promoter [55] and either a Cas12a Full Direct Repeat, Mature Direct Repeat or a truncated tRNA fused to Mature Direct Repeat together with a ccdB-CmR cassette flanked by BbsI sites and introduced in entry clones by Gibson assembly [56]. This resulted in the following entry clones: pCG264_pGG-A-TaU3-full-DR-LbCas12a-BsbI-ccdB-BsbI-B (p2830), pCG265_pGG-A-TaU3-mature-DR-LbCas12a-BsbI-ccdB-BsbI-B (p2831) and pCG266_pGG-A-TaU3-tRNA-mature-DR-LbCas12a-BsbI-ccdB-BsbI-B (p2832). New crRNAs were cloned via Golden Gate with annealed oligos as described above. To facilitate the cloning of multiplex TaU3:tRNA-mature-DR-crRNA vectors, we created a set of entry clones with A-B (PSB:17_79), B-C (PSB:17_80), C-D (PSB:17_81), D-E (PSB:18_01), E-F (PSB:18_02) and F-G (PSB:18_03) GreenGate overhangs by Gibson assembly of TaU3, ccdB-Cm PCR products and stitching oligos as previously described[57]. All primers are described in Additional file 7.

#### Base editors

Base editor expression vectors were assembled following the general architecture: pGG-A-promoter-B, pGG-B-deaminase-C, pGG-C-nuclease-D, pGG-D-NLS-E, pGG-E-terminator-F and pGG-F-linker-G into the destination vectors p35S-mCherry-NLS_CmR-ccdB (p02245) or pGG-AG-KmR (p00343) via Golden Gate with BsaI.

#### Fluorescent reporters

The FLARE system was used for CBE fluorescent reporters [24]. We introduced a T63S substitution into the BFP sequence to add a Cas12a PAM site (p02238). For the ABE-nCas9 fluorescent reporter (p02662), we substituted the Q70 residue of GFP with a stop codon (TAG).

For the Cas12a-ABE reporter 1 (p02236), we substituted the R110 residue of GFP with a stop codon (TGA). For the Cas12a-ABE reporter 2 (p02877), we substituted the Q70 residue of GFP with a stop codon (TAG) and added a PAM site by introducing R74K mutation as previously described [19]. All site-directed mutagenesis was performed with Gibson assembly and expression vectors were assembled by Golden Gate with BsaI.

#### Vector isolation for protoplast transfection

Plasmids were purified using the plasmid Midiprep kit (D4200, Zymo Research) and Maxi kit (12163, QIAGEN) according to manufacturer protocols. Vector concentration was measured using nanodrop (Isogen LifeSciences).

### Plant growth conditions

Wheat seeds (Fielder) were processed by successive washes with sterilized water for 3 min, isopropanol for 45 sec, sterile water for 3 min and 6% sodium hypochlorite (Chem-lab nv) for 10 min. Sterilized seeds were washed six times with sterile water in a laminar flow cabinet and sown on sterile growth media containing 1/2 MS pH 5.7 (Duchefa Biochemie, M0221.0050), 2.5 mM MES (M1503.0100, Duchefa Biochemie) and 0.5% plant tissue agar (NCM0250A, NEOGEN). Two seeds were sown per sterile, 175 ml cylindrical container (960162, Greiner Bio-one) and stratified for 3 days at 4°C in the dark. Plants were grown under SpectraluxPlus NL 36 W/ 840 Plus (Radium Lampenwerk) fluorescent bulbs under long days (16h light/8h dark) at 25°C.

B104 maize seeds were sown directly on Jiffy substrate (No. 32170138, Jiffy Products International). Seed germination was performed in long day (16h light/8h dark) conditions at 25°C, 55% relative humidity for 5 days under light provided by high-pressure sodium vapor (RNP-T/LR/400W/S/230/E40, Radium) and metal halide lamps with quartz burners (HRI-BT/400W/D230/E40, Radium). Seedlings were transferred to the dark for 8 days prior to protoplast isolation.

### Protoplast isolation and transfection

Wheat protoplast isolation was adapted from [58]. Wheat leaves were harvested 7 or 8 days after germination (DAG). Approximately 40-50 second leaves were cut into latitudinal 0.5-1 mm strips with a sharp razor blade and leaf strips were incubated in 0.6 M D-mannitol (M1902, Sigma-Aldrich) for 10 min in the dark. The mannitol was removed and 25 ml of cell wall enzyme solution (20 mM MES, 1.5% cellulase R10 (C8001.0010), 0.75% macerozyme R10 (M8002.0010), 0.6 M D-mannitol and 10 mM KCl, 0.1% BSA and 10 mM CaCl_2_) was added to protoplasts for 8 hours incubation in the dark at 25°C with shaking at 40 rpm. After enzymatic digestion, 25 ml of W5 solution (2 mM MES pH 5.7, 154 mM NaCl, 125 mM CaCl_2_ 0,5 mM KCl) was added to release the protoplasts. Protoplasts were collected by filtering the mixture through a sterile 40 μm cell strainer (#431750, Corning) and centrifuging at 80*g* (slow acceleration and brake) for 3 min at room temperature. The supernatant was discarded and protoplasts were resuspended in 6 ml W5 solution and incubated on ice for 30 min. Protoplast yield was determined using a Neubauer chamber before adding MMG_Ta_ solution (4 mM MES pH 5.7, 0.4 M mannitol, 15 mM MgCl2) onto the cell pellet to reach a concentration of 1 × 10^6^ cells ml^−1^.

Protoplasts were incubated on ice for ~30 min before transfection. 12 μg of total plasmid DNA was added to MMG_Ta_ to a total volume of 20 μl in 1 ml strip tubes (TN0946-08B, National Scientific Supply Co). 100 μl of protoplasts (10^5^ cells) and 110 μl of PEG solution (0.2 M mannitol, 100 mM CaCl_2_), 40% PEG 4000 (81240, Sigma) were added using a multichannel pipette to DNA and immediately mixed by slowly inverting the strip. For individual strips, 8 transfections were processed in parallel. Protoplasts were incubated for 15–20 min and W5 solution was added to stop the transfection. After centrifuging at 80*g* (slow acceleration and brake) for 3 min, the supernatant was discarded and the protoplast pellet resuspended in 1 ml of W5 solution. Cells were transferred in 24-well plates (734-2325, VWR) and incubated in the dark at 25°C for 42 to 46 hours.

Maize protoplast isolation was adapted from [59,60]. Etiolated maize leaves were harvested at 12 or 13 DAG. The middle part of the second or third leaf was cut into 0.5 mm strips. Strips were then infiltrated with 25 ml cell wall enzyme solution (0.6 M D-mannitol, 10 mM MES, 1.5% cellulose, 0.3% Macerozyme R10, 0.1% BSA and 1 mM CaCl_2_) using vacuum (50 mmMg) for 30 minutes in the dark and then incubated for 2 hours at 25°C with shaking (40 rpm). The solution containing the protoplasts was filtered using a sterile 40 μm cell strainer (Corning) and collected by centrifuging at 100*g* (slow acceleration and brake) for 3 min. The supernatant was removed and protoplasts washed with ice-cold 0.6 M D-mannitol by centrifugation at 100*g* (slow acceleration and brake) for 2 min. Cells were resuspended in 5 ml of 0.6 M D-mannitol and incubated in the dark for 30 min. Protoplasts were resuspended in MMG_Zm_ solution (0.6 M mannitol, 15 mM MgCl_2_, 4 mM MES) and counted using a Neubauer chamber and adjusted to a concentration of 1 × 10^6^ cells ml^−1^.

20 μg of total plasmid DNA was added to MMG_Zm_ to a total volume of 20 μl in 1 ml strip tubes. 100 μl of protoplasts (10^5^ cells) and 110 μl of PEG (0.2 M mannitol, 100 mM CaCl_2_, 40% PEG 4000) solution were added using a multichannel pipette to DNA and immediately mixed by inverting the strip. For individual strips, 8 transfections were processed in parallel. Cells were then incubated for 10–15 min in the dark and W5 solution was added to stop the transfection. After centrifugation at 100*g* (slow acceleration and brake) for 2 min, supernatant was discarded and the protoplast pellet resuspended in 1 ml of W5 solution. Cells were then transferred using tips with wide bore in 24-well plates (VWR) and incubated in the dark at 25°C with shaking (20 rpm) for 2 days.

### High content image analysis

Two days after transfection, 50 μl of protoplasts were transferred to 96-well Cell carrier Ultra plates (6055302, PerkinElmer) and imaged with the Opera Phenix® High Content Screening System (PerkinElmer). Image acquisition was performed using a 20× water immersion objective in confocal mode, taking 7 Z-planes and 9 fields of view per well and covering 4 image channels: brightfield, Chlorophyl, GFP and mCherry. Raw images were transferred to the Columbus™ Image Data Storage and Analysis system for automated image processing and quantification.

After flatfield correction and smoothing of the chlorophyll channel, single wheat cells were segmented and selected as protoplasts based on roundness. mCherry and GFP signals were used to identify nuclei and exclude non-transformed protoplasts based on the absence of nuclear mCherry signal. The mCherry and GFP intensities in the nuclei of transformed protoplasts were used to identify and quantify the GFP expressing transformed protoplasts.

For maize, the chlorophyll channel could not be used for cell segmentation as the plants were etiolated. The analysis focused directly on transformed protoplast nuclei, segmenting based on mCherry and GFP channels. The same analysis as described above was used to identify and quantify the GFP expressing transformed nuclei.

Results were exported as a table and all calculations and image processing were performed on an in-house cluster (VIB). The time required from the start of imaging to obtaining processed results takes 3-4 hours for a 96-well plate. Codes for wheat and maize analysis workflows are available in Additional files 4-5.

### FACS

Images were captured using a BD Biosciences FACS imaging enabled prototype cell sorter that is equipped with an optical module allowing multicolor fluorescence imaging of fast flowing cells in a stream enabled by BD CellView™ Image Technology based on fluorescence imaging using radiofrequency-tagged emission (FIRE)[61,62].

Two days after transfection, 500 μl of protoplast solution was used for sorting. Gating strategies for GFP were first established on cells expressing pZmUBI-GFP-NLS (p02243) and similar settings were used for all experiments in wheat and maize. A quality check was conducted by running the sorted cell fraction on the instrument and imaged using the imaging system integrated in the FACS instrument. For both wheat and maize, a 130 μm nozzle was used and 1,000 to 5,000 cells were sorted into 1.5 ml Eppendorf tubes containing 10 μl of dilution buffer from the Phire Tissue Direct PCR Master Mix kit (F160L, Thermo Fisher Scientific).

### Genotyping and NGS analysis

For genotyping individual wheat transformed plants, DNA was isolated according to the Edwards method [63]. A piece of leaf (0.5-1 cm), harvested in 1 ml tubes on 96-well plate (732-3716, VWR) and flash frozen in liquid nitrogen. Two metal beads (3 mm) were added and tissue was ground to a powder by shaking the plate at 20 Hz for 1 minute (Mixer Mill MM 400, Retsch). 400 μl of extraction buffer (100 mM Tris-HCl pH 8.0, 500 mM NaCl, 50 mM EDTA, 0.7% SDS) was added to individual samples and incubated for 30 min at 60°C. Samples were centrifuged and 300 μl of the supernatant was mixed with 300 μl of isopropanol for DNA precipitation. Samples were then centrifuged and supernatant removed. The pellet was washed with 70% Ethanol, dried at room temperature, and dissolved in 100 μl of 10 mM Tris-HCl pH 8.0.

For sorted material, 2 μl of the solution containing sorted cells was used as template in a 20 μl total reaction volume for amplicon PCR using the Phire Plant Direct PCR Kit, according to manufacturer’s recommendations. All raw gels can be found in Appendix (Additional file 1).

For nCas9-BE, base editing efficiencies were measured by Sanger sequencing. 500-700 bp amplicons were designed to amplify targets (Additional file 7-8). 5 μl of the Phire PCR reaction was verified on a 1% agarose gel (Ultra-Pure agarose, # 16500-500, Invitrogen), with a 1kb bench top ladder (G754A, Promega). 15 μl was purified using 1.8× HighPrep™ PCR beads (AC-60500, MAGBIO Genomics), eluted in 35 μl 10 mM Tris-HCl pH 8.0 and sent for Sanger sequencing (Mix2Seq, Eurofins). Chromatograms were processed using EditR to obtain base editing efficiencies[64]. Statistical tests for normality and comparative analysis were conducted with GraphPad Prism 9.0.0 or VassarStats online tool (http://vassarstats.net/). For dCas12-BE and nuclease-active LbCas12a, base editing and indel efficiencies were measured using NGS. 210-260 bp amplicons were designed to amplify targets. 6-nt indices[65] were added to forward and reverse primers for pooling and demultiplexing amplicon reads after sequencing (Additional file 7). 5 μl of the Phire PCR reaction was verified on a 2% agarose gel with a low molecular weight ladder (N3233S, NEB). 15 μl of PCR products were pooled and purified using PCR Purification Kit (D4013, Zymo Research Co.). Depending on the amplicon, an extra gel purification was conducted (D4002, Zymo Research) to specifically isolate the PCR band of the target site. The DNA concentration was measured with Qubit (Invitrogen) according to manufacturer’s protocol and adjusted to 2 ng μl^−1^. Paired end sequencing was performed with Eurofins NGSelect amplicons (5M reads 2×150bp). Reads were demultiplexed using Je-demultiplex [66] and individual fastq files were obtained as previously described [67] using a Galaxy workflow (https://usegalaxy.be). Demultiplexing statistics can be found in Additional file 3.

Base editing was calculated using CRISPResso2Pooled or CRISPRessoBatch [68]. Editing window and read quality were defined as follows: cleavage offset was set to −1, quantification window size to 10, quantification window center to −12 and minimum average read quality to 30. Indels were calculated using CRISPResso2Pooled or CRISPRessoBatch with the following settings to define Cas12a cutting site and read quality: cleavage offset was set to −4 and minimum average read quality to 30. All amplicons and gRNA sequences can be found in Additional file 8.

### Stable wheat transformation

Immature embryos 2-3 mm in size were isolated from sterilized ears of wheat cv. Fielder and bombarded using the PDS-1000/He particle delivery system (Bio-Rad) essentially as previously described [69] using the following particle bombardment parameters: diameter gold particles, 0.6 μm; target distance, 6 cm; bombardment pressure, 7.584 kPa; gap distance, 8-10 mm; microcarrier flight distance, 10 mm; vacuum within the bombardment chamber, 27.5” Hg. For each shot approximately 150 μg of gold particles carrying 570 ng of plasmid DNA were delivered.

The applied plasmid DNA was a mixture of the Cas12a-ABE vectors pCG392 or pCG434, pCG406 and pCG408 (gRNAs) and pBAY02032 (selectable marker). The vector pBAY02032 contains an eGFP-BAR fusion gene under control of the 35S promoter. The further culture of the bombarded immature embryos was essentially conducted as previously described [70]. Bombarded immature embryos were transferred to non-selective WLS callus induction medium for about one week, then moved to WLS with 5 mg L^−1^ phosphinothricin (PPT) for a first selection round of about 3 weeks followed by a second selection round on WLS with 10 mg L^−1^ PPT for another 3 weeks. PPT resistant calli were selected and transferred to shoot regeneration medium with 5 mg L^−1^ PPT.

## Additional file 1: information

Additional file 1: Additional file 1: figures. **Fig. S1.** Establishment of high content analysis pipeline. **Fig.S2.** ITER validation for Cas9 base editors using GFP reporters. **Fig. S3.** ITER validation for Cas9 base editors at endogenous sites. **Fig. S4**. Iterative optimization of Cas12a-ABEs using ITER. **Fig. S5.** Cas12a-ABEs efficiency at endogenous sites in protoplasts. **Fig. S6.** Cas12a-ABE efficiency in stable wheat transformants. **Fig. S7.** Cas12a-CBEs and LbCas12a efficiency at endogenous sites in protoplasts. **Appendix.** Raw gel pictures.

Additional file 2: Raw data file

Additional file 3: Statistics NGS

Additional file 4: Workflow OPERA wheat

Additional file 5: Workflow OPERA maize

Additional file 6: Plasmid list

Additional file 7: Primer list

Additional file 8: Target sites list

Additional file 9: NGS summaries

## Acknowledgments

We thank Hilde Nelissen and Jie Zhang for assistance with maize protoplast transfections. We thank Kristel D’Hont, Martine Bossut, Greet De Backer, Anne-Marie De Loof, Anouk Pennewaert, Bernadette Saey, Chantal Vanderstraeten and Sigrid Vanhoutte for the production and analysis of stably transformed wheat plants. We thank the VIB Technology Watch fund for assistance with DNA synthesis.

## Authors’ contributions

C.G, A.P.F, F.M, T.B.J designed the study. F.M, T.B.J supervised the study. C.G, A.P.F, CaG, J.D.S and D.D.V conducted protoplast transfections. C.G, A.P.F conducted all other experimental assays and analyses. V.G set up the OPERA image analysis pipeline. V.G, A.D and D.A provided technical assistance with the OPERA system. J.V.D, G.V.I set up FACS sorting pipeline and provided technical assistance. C.G, M.F, W.D designed targets and primers. C.G and W.D analyzed NGS data. K.D.H, D.D.V conducted stable wheat transformations and preliminary genotyping. C.G, A.P.F, F.M, T.B.J prepared figures and wrote the manuscript with contribution from all co-authors.

## Availability of data and materials

Plasmids and vector maps (.gb files) are available from https://gatewayvectors.vib.be/.

Amplicon NGS files were deposited at GEO (https://www.ncbi.nlm.nih.gov/geo/) under the accession numbers: GSE200450. A summary file can be found in Additional file 9.

