## Additional file 1 for "Systematic optimization of Cas12a base editors in wheat and maize using the ITER platform"

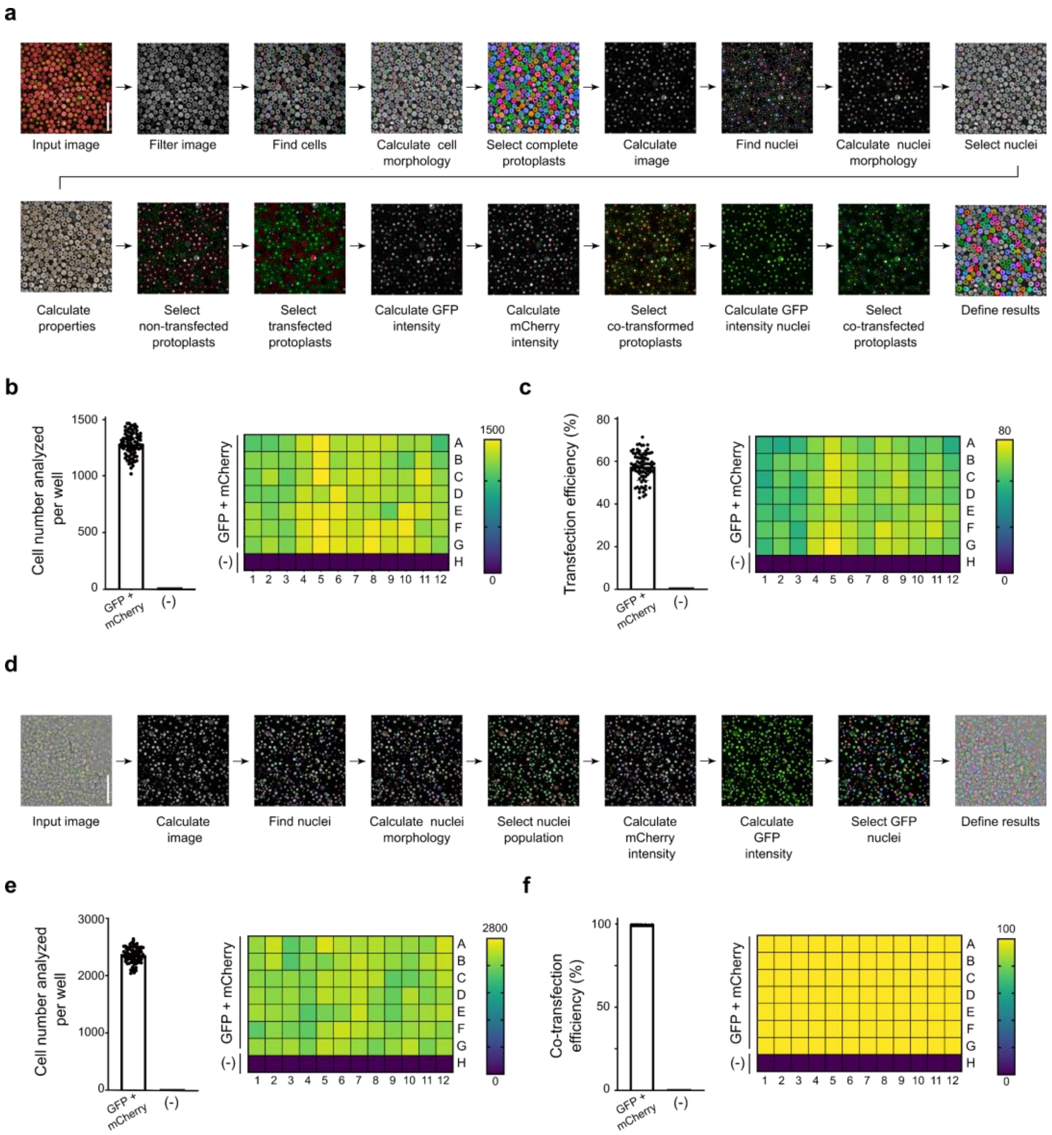


**Fig. S1: Establishment of high content analysis pipeline.**

**a** Representative images from image segmentation and quantification workflow for wheat protoplasts on the OPERA® High Content Screening System. **b** Cell number at individual wells after protoplast transfection as measured by the total number of mCherry positive wheat cells. **c** Transfection efficiency at individual wells as measured by mCherry positive wheat cells over the total number of segmented cells. **d** Representative images from image segmentation and quantification workflow for maize protoplasts on high content imaging platform. **e** Cell number at individual wells after protoplast transfection as measured by the total number of mCherry positive maize cells. **f** Co-transfection efficiency as measured by the number of GFP over the number of mCherry positive cells in maize. Negative controls (-) do not contain vector DNA (b,c,e,f). The grid shows the 96-well plate positions and gradient color coding represent number of cell analyzed (b,e), transfection efficiencies (c) or co-transfection efficiency (f). Scale bars in a,d: 200 µm

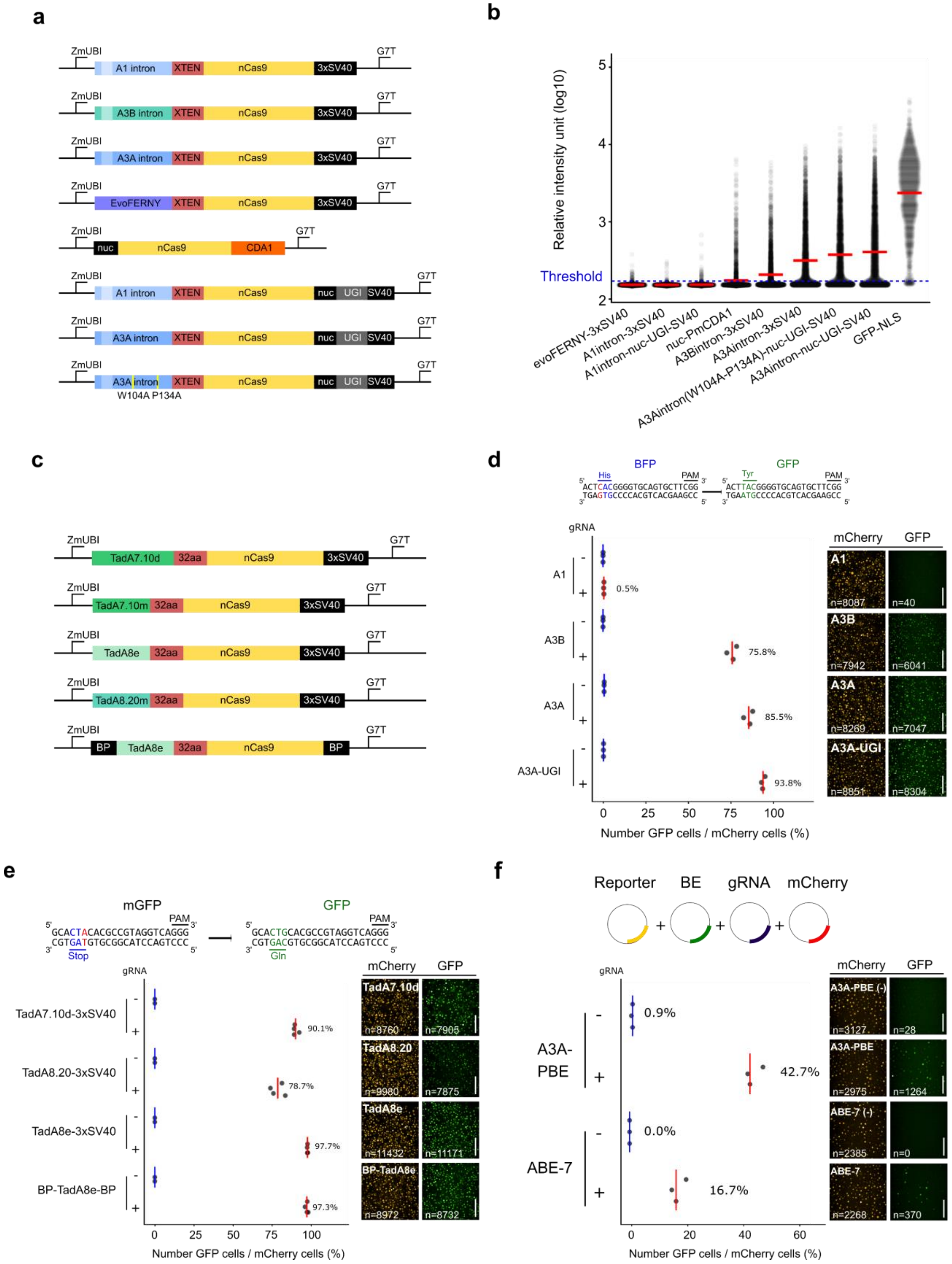
a
ZmUBI
G7T
A1 intron
A3B intron
A3A intron
EvoFERNY
XTEN
nCas9
3xSV40
nuc
CDA1
UGI
SV40
W104A P134A
b
Relative intensity unit (log10)
Threshold
evoFERNY-3xSV40
A1intron-3xSV40
A1intron-nuc-UGI-SV40
nuc-PmCDA1
A3Bintron-3xSV40
A3Aintron-3xSV40
A3Aintron(W104A-P134A)-nuc-UGI-SV40
A3Aintron-nuc-UGI-SV40
GFP-NLS
c
TadA7.10d
TadA7.10m
TadA8e
TadA8.20m
32aa
BP
d
BFP
GFP
His
Tyr
PAM
ACTCACGGGGTGCAGTGCTTCGG
TGAGTGCCCCACGTCACGAAGCC
ACTTACGGGGTGCAGTGCTTCGG
TGAATGCCCCACGTCACGAAGCC
gRNA
A1
A3B
A3A
A3A-UGI
0.5%
75.8%
85.5%
93.8%
mCherry
GFP
n=8087
n=40
n=7942
n=6041
n=8269
n=7047
n=8851
n=8304
Number GFP cells / mCherry cells (%)
e
mGFP
GCACTACACGCCGTAGGTCAGGG
CGTGATGTGCGGCATCCAGTCCC
Stop
GCACTGCACGCCGTAGGTCAGGG
CGTGACGTGCGGCATCCAGTCCC
Gln
TadA7.10d-3xSV40
TadA8.20-3xSV40
TadA8e-3xSV40
BP-TadA8e-BP
90.1%
78.7%
97.7%
97.3%
TadA7.10d
TadA8.20
TadA8e
BP-TadA8e
n=8760
n=7905
n=9980
n=7875
n=11432
n=11171
n=8972
n=8732
f
Reporter
BE
gRNA
mCherry
A3A-PBE
ABE-7
0.9%
42.7%
0.0%
16.7%
A3A-PBE (-)
ABE-7 (-)
n=3127
n=28
n=2975
n=1264
n=2385
n=0
n=2268
n=370

**Fig. S2: ITER validation for Cas9 base editors using GFP reporters**

**a** Schematic representation of Cas9-CBE architectures tested in this study. **b** Relative GFP intensity of individual cells transfected with nCas9 CBEs. The GFP-NLS positive control was transfected with pZmUBI:GFP-NLS. Red bars represent mean. **c**, Schematic representation of Cas9-ABE architectures tested in this study. **d-e** Base editing efficiencies for Cas9-CBEs (d) and Cas9-ABEs (e) as measured by the rate GFP conversion in maize cells. **f** Base editing efficiency for A3A-PBE [27] and ABE-7 [28] as determined by the rate of GFP conversion in wheat cells. Editing rates were calculated from three or four independent biological replicates that are depicted as dots. Red and blue lines represent mean efficiencies with or without gRNAs, respectively. Representative pictures of protoplasts showing mCherry or GFP fluorescence. n indicates the total number of cells measured after image segmentation in the mCherry and the GFP channel (d-f). Scale bars in d-f: 200 µm

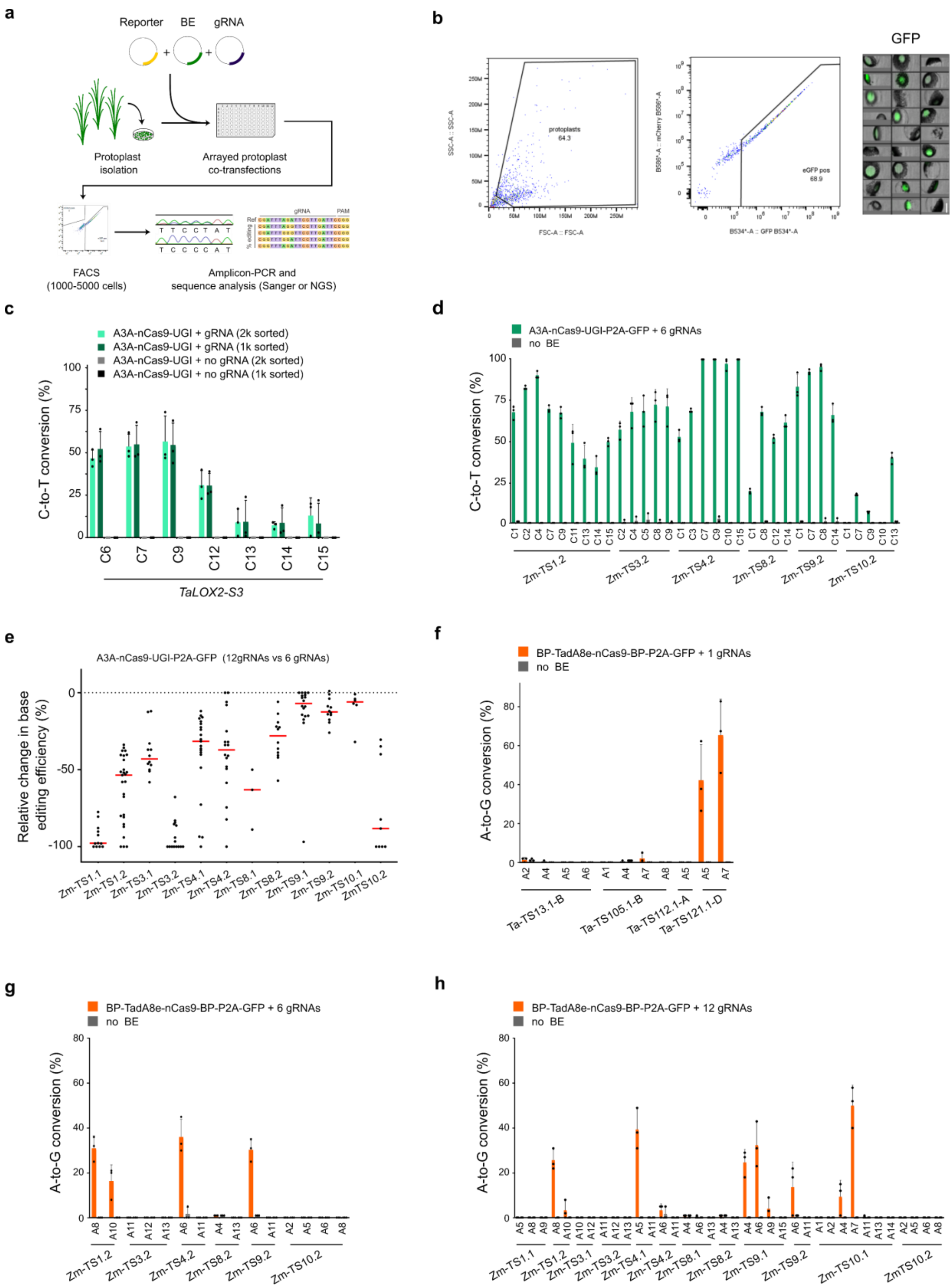

a
Reporter
BE
gRNA
Protoplast isolation
Arrayed protoplast co-transfections
FACS (1000-5000 cells)
Amplicon-PCR and sequence analysis (Sanger or NGS)
b
GFP
protoplasts 64.3
eGFP pos 68.9
c
A3A-nCas9-UGI + gRNA (2k sorted)
A3A-nCas9-UGI + gRNA (1k sorted)
A3A-nCas9-UGI + no gRNA (2k sorted)
A3A-nCas9-UGI + no gRNA (1k sorted)
C-to-T conversion (%)
TaLOX2-S3
d
A3A-nCas9-UGI-P2A-GFP + 6 gRNAs
no BE
C-to-T conversion (%)
Zm-TS1.2
Zm-TS3.2
Zm-TS4.2
Zm-TS8.2
Zm-TS9.2
Zm-TS10.2
e
A3A-nCas9-UGI-P2A-GFP (12gRNAs vs 6 gRNAs)
Relative change in base editing efficiency (%)
Zm-TS1.1
Zm-TS1.2
Zm-TS3.1
Zm-TS3.2
Zm-TS4.1
Zm-TS4.2
Zm-TS8.1
Zm-TS8.2
Zm-TS9.1
Zm-TS9.2
Zm-TS10.1
ZmTS10.2
f
BP-TadA8e-nCas9-BP-P2A-GFP + 1 gRNAs
no BE
A-to-G conversion (%)
Ta-TS13.1-B
Ta-TS105.1-B
Ta-TS112.1-A
Ta-TS121.1-D
g
BP-TadA8e-nCas9-BP-P2A-GFP + 6 gRNAs
no BE
A-to-G conversion (%)
h
BP-TadA8e-nCas9-BP-P2A-GFP + 12 gRNAs
no BE
A-to-G conversion (%)

**Fig. S3: ITER validation for Cas9 base editors at endogenous sites.**

**a** Schematic representation of Fluorescence-Activated Cell Sorting (FACS) and genotyping workflows. Vectors are co-transfected in protoplasts and sorted on GFP signal directly in dilution buffer. Solution containing lysed protoplasts is used as template for PCR and resulting amplicons are used for Sanger sequencing or NGS. **b** Gating strategy used for sorting protoplasts expressing GFP. Pictures show intact wheat cells expressing nuclear GFP after sorting. Scale bar**:** 20 µm. **c** Simplex base editing efficiency of A3A-nCas9-UGI at *TaLOX2-S3* (c) as measured by Sanger sequencing. One wheat homeolog was analyzed. Transfection of the CBE without the gRNA was used as a negative control. **d** Multiplex base editing efficiency of A3A-nCas9-UGI at six maize targets. Percentages indicate the proportion of C-to-T base conversion. Transfection of the CBE without the gRNA was used as a negative control. The x-axis indicates targeted Cytosines at different positions along the protospacer, with the PAM-distal base being position 1 (c-e). **e** Relative change in multiplex base editing efficiency between 6 and 12 gRNAs in maize protoplasts. **f** Simplex base editing efficiency of BP-TadA8e-nCas9-BP at four wheat targets. **g-h** Multiplex base editing of BP-TadA8e-nCas9-BP at 6 (g) or 12 (h) maize targets. The x-axis indicates targeted Adenine at different positions along the protospacer, with the PAM-distal base being position 1 (g-i). Editing rates were calculated from three independent biological replicates that are depicted as dots (c-i). Transfection of the cells with a GFP expressing vector (no BE) was used as a negative control (e-i).

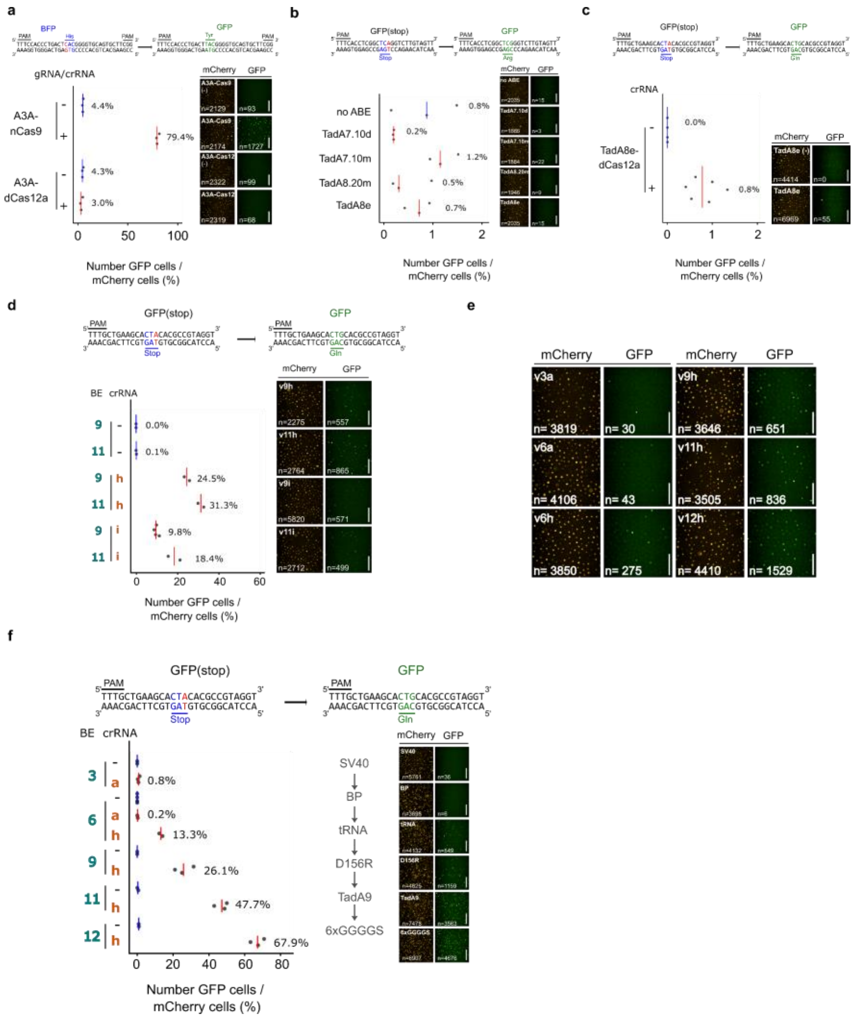


**Fig. S4: Iterative optimization of Cas12a-ABEs using ITER**

**a** Base editing efficiencies for A3A-nCas9 or A3A-dCas12a in wheat as measured by the rate of BFP to GFP conversion. **b** Base editing efficiencies for Cas12a-ABEs as measured by the rate of GFP recovery (stop to Arg) in wheat cells. **c** Base editing efficiency for TadA8e-dCas12a as measured by the rate of GFP recovery (stop to Gln) in wheat cells. **d** Base editing efficiency for Cas12a-ABEs as measured by the rate of GFP recovery (stop to Gln) in wheat cells. **e** Representative pictures showing mCherry and GFP positive cells measured during the comparative analysis of key Cas12a-BE architectures in Fig2. c''. **f** Comparative analysis of key Cas12a-ABE architectures obtained along the optimization path as measured by the rate of GFP recovery (from stop to Gln) in maize cells. Components leading to increased activity are shown at the right of the panel. Representative pictures show fluorescence measured for mCherry and GFP cells and the number of cells analyzed. Editing rates were calculated from two to four independent biological replicates that are depicted as dots. Scale bars in a-f: 200 µm

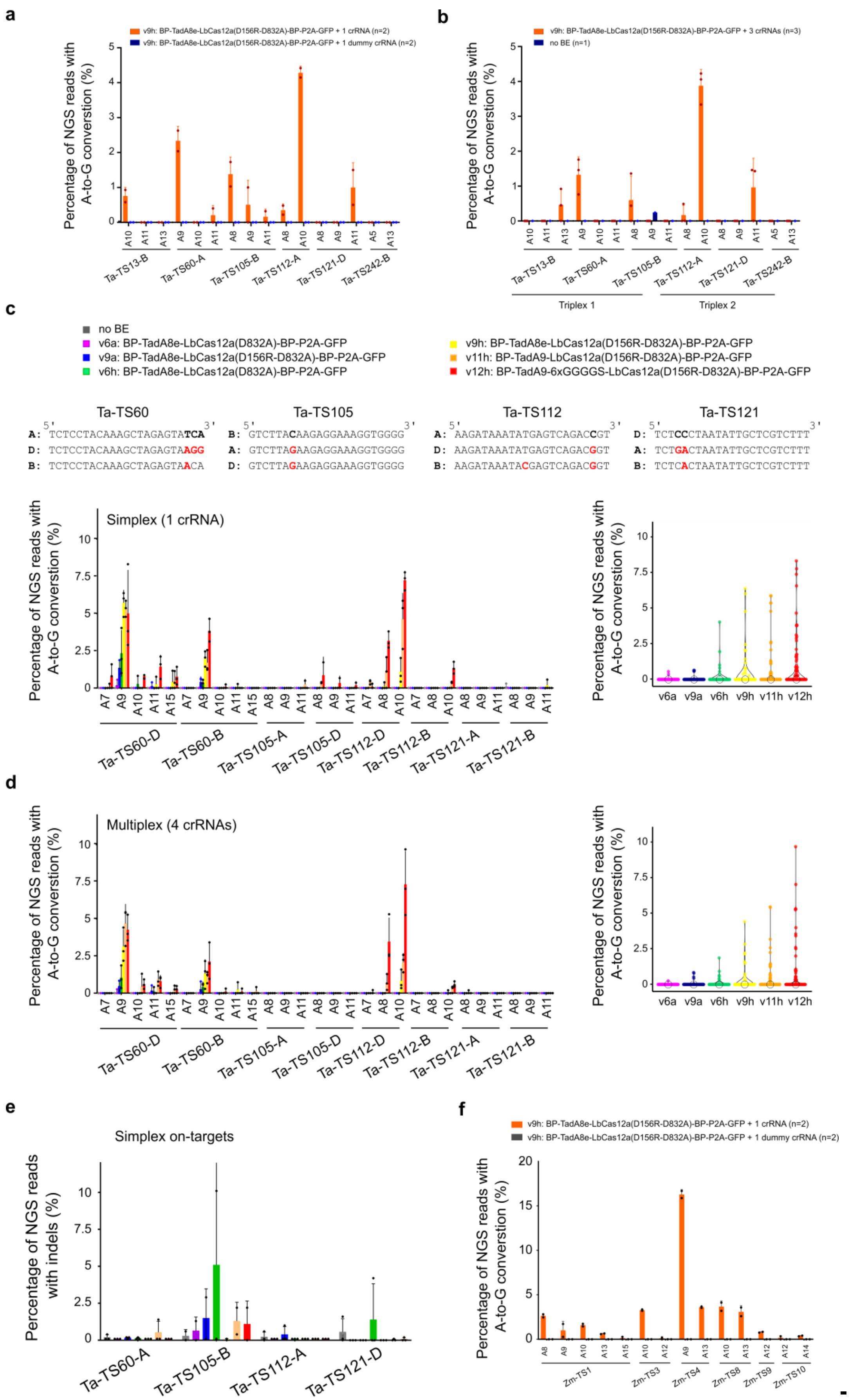

a
v9h: BP-TadA8e-LbCas12a(D156R-D832A)-BP-P2A-GFP + 1 crRNA (n=2)
v9h: BP-TadA8e-LbCas12a(D156R-D832A)-BP-P2A-GFP + 1 dummy crRNA (n=2)
Percentage of NGS reads with A-to-G conversion (%)
Ta-TS13-B
Ta-TS60-A
Ta-TS105-B
Ta-TS112-A
Ta-TS121-D
Ta-TS242-B
b
v9h: BP-TadA8e-LbCas12a(D156R-D832A)-BP-P2A-GFP + 3 crRNAs (n=3)
no BE (n=1)
Percentage of NGS reads with A-to-G conversion (%)
Triplex 1
Triplex 2
c
no BE
v6a: BP-TadA8e-LbCas12a(D832A)-BP-P2A-GFP
v9a: BP-TadA8e-LbCas12a(D156R-D832A)-BP-P2A-GFP
v6h: BP-TadA8e-LbCas12a(D832A)-BP-P2A-GFP
v9h: BP-TadA8e-LbCas12a(D156R-D832A)-BP-P2A-GFP
v11h: BP-TadA9-LbCas12a(D156R-D832A)-BP-P2A-GFP
v12h: BP-TadA9-6xGGGGS-LbCas12a(D156R-D832A)-BP-P2A-GFP
Ta-TS60
A: TCTCCTACAAAGCTAGAGTATCA
D: TCTCCTACAAAGCTAGAGTAAGG
B: TCTCCTACAAAGCTAGAGTAACA
Ta-TS105
B: GTCTTACAAGAGGAAAGGTGGGG
A: GTCTTAGAAGAGGAAAGGTGGGG
D: GTCTTAGAAGAGGAAAGGTGGGG
Ta-TS112
A: AAGATAAATATGAGTCAGACCGT
D: AAGATAAATATGAGTCAGACGGT
B: AAGATAAATACGAGTCAGACGGT
Ta-TS121
D: TCTCCCTAATATTGCTCGTCTTT
A: TCTGACTAATATTGCTCGTCTTT
B: TCTCACTAATATTGCTCGTCTTT
Simplex (1 crRNA)
Percentage of NGS reads with A-to-G conversion (%)
Ta-TS60-D
Ta-TS60-B
Ta-TS105-A
Ta-TS105-D
Ta-TS112-D
Ta-TS112-B
Ta-TS121-A
Ta-TS121-B
v6a v9a v6h v9h v11h v12h
d
Multiplex (4 crRNAs)
Percentage of NGS reads with A-to-G conversion (%)
e
Simplex on-targets
Percentage of NGS reads with indels (%)
Ta-TS60-A
Ta-TS105-B
Ta-TS112-A
Ta-TS121-D
f
v9h: BP-TadA8e-LbCas12a(D156R-D832A)-BP-P2A-GFP + 1 crRNA (n=2)
v9h: BP-TadA8e-LbCas12a(D156R-D832A)-BP-P2A-GFP + 1 dummy crRNA (n=2)
Percentage of NGS reads with A-to-G conversion (%)
Zm-TS1
Zm-TS3
Zm-TS4
Zm-TS8
Zm-TS9
Zm-TS10

**Fig. S5: Cas12a-ABEs efficiency at endogenous sites in protoplasts**

**a-b** Base editing efficiencies of TadA8e-dCas12a (v9h) in simplex (a) or multiplex (b) as measured by the proportion of NGS reads converted from A-to-G at six wheat targets. **c-d** Base editing efficiencies of six Cas12a-ABE configurations in simplex or multiplex at four wheat off-targets located on homoeologous genes as measured by the proportion of NGS reads converted from A-to-G. DNA sequences on the top row are the on-targets, red bases indicate mismatches with the crRNA at off-target homeologs. **e** Indel rate measured at wheat on-targets for six Cas12a-ABE configurations. Color code for Cas12a-ABE architectures is same as (c). **f** Simplex Base editing efficiencies of TadA8e-dCas12a (v9h) in maize as measured by the proportion of reads converted from A-to-G at four targets. Editing rates were calculated from two or three independent biological replicates.

a

| | | TaTS60-D | | | | | TaTS60-B | | | | | TaTS60-A | | | | |
|---|---|---|---|---|---|---|---|---|---|---|---|---|---|---|---|---|
| | | A7 | A9 | A10 | A11 | A15 | A7 | A9 | A10 | A11 | A15 | A7 | A9 | A10 | A11 | A15 |
| v9h | Mutated (%) | 6.5 | 31.8 | 2.6 | 0.6 | 0.0 | 1.3 | 16.9 | 0.0 | 0.0 | 0.0 | 3.8 | 16.1 | 0.6 | 0.0 | 0.6 |
| v11h | Mutated (%) | 15.8** | 40.4 | 13.1*** | 7.7** | 1.6 | 8.2** | 26.5* | 0.0 | 1.1 | 0.5 | 11.5** | 34.6*** | 3.8 | 3.3* | 0.5 |

| | | TaTS112-D | | | | TaTS112-B | | | | TaTS112-A | | | |
|---|---|---|---|---|---|---|---|---|---|---|---|---|---|
| | | A6 | A7 | A8 | A10 | A6 | A7 | A8 | A10 | A6 | A7 | A8 | A10 |
| v9h | Mutated (%) | 1.3 | 0.0 | 1.3 | 24.8 | 0.0 | 0.0 | 0.0 | 1.9 | 1.9 | 0.0 | 2.6 | 34.0 |
| v11h | Mutated (%) | 3.8 | 1.1 | 4.4 | 39.6** | 0.0 | 0.0 | 0.0 | 6.6* | 4.4 | 0.5 | 12.1*** | 46.7* |

0 50

Percentage of T0 lines with A-to-G conversion (%)

b

| | | TaTS60-D | | | | | TaTS60-B | | | | | TaTS60-A | | | | |
|---|---|---|---|---|---|---|---|---|---|---|---|---|---|---|---|---|
| | | A7 | A9 | A10 | A11 | A15 | A7 | A9 | A10 | A11 | A15 | A7 | A9 | A10 | A11 | A15 |
| v9h | HZ (%) | 6.5 | 24.0 | 2.6 | 0.0 | 0.0 | 1.3 | 15.6 | 0.0 | 0.0 | 0.0 | 3.2 | 12.9 | 0.6 | 0.0 | 0.6 |
| | HM (%) | 0.0 | 7.8 | 0.0 | 0.6 | 0.0 | 0.0 | 1.3 | 0.0 | 0.0 | 0.0 | 0.6 | 3.2 | 0.0 | 0.0 | 0.0 |
| v11h | HZ (%) | 14.8* | 30.6 | 13.1* | 7.1*** | 1.6 | 8.2** | 22.7 | 0.0 | 1.1 | 0.5 | 10.4* | 28.0*** | 3.8 | 3.3* | 0.5 |
| | HM (%) | 1.1 | 9.8 | 0.0 | 0.5 | 0.0 | 0.0 | 3.9 | 0.0 | 0.0 | 0.0 | 1.1 | 6.6 | 0.0 | 0.0 | 0.0 |

| | | TaTS112-D | | | | TaTS112-B | | | | TaTS112-A | | | |
|---|---|---|---|---|---|---|---|---|---|---|---|---|---|
| | | A6 | A7 | A8 | A10 | A6 | A7 | A8 | A10 | A6 | A7 | A8 | A10 |
| v9h | HZ (%) | 1.3 | 0.0 | 1.3 | 21.0 | 0.0 | 0.0 | 0.0 | 1.9 | 1.9 | 0.0 | 2.6 | 23.7 |
| | HM (%) | 0.0 | 0.0 | 0.0 | 3.8 | 0.0 | 0.0 | 0.0 | 0.0 | 0.0 | 0.0 | 0.0 | 10.3 |
| v11h | HZ (%) | 3.8 | 1.1 | 4.4 | 28.0 | 0.0 | 0.0 | 0.0 | 6.6* | 4.4 | 0.5 | 11.5** | 28.6 |
| | HM (%) | 0.0 | 0.0 | 0.0 | 11.5** | 0.0 | 0.0 | 0.0 | 0.0 | 0.0 | 0.0 | 0.5 | 18.1* |

0 50

Percentage of T0 lines with A-to-G conversion (%)

### Fig. S6: Cas12a-ABE efficiency in stable wheat transformants

**a-b** Frequency of individual T0 wheat plants carrying heterozygous or homozygous A-to-G conversion at individual positions of the targets. Combined heterozygous (HZ) and homozygous (HM) frequencies are shown in a., individual frequencies in b. Values are depicted for TaTS60 and TaTS112 at the three subgenomes (n=154 for v9h and n=184 for v11h). Asterisks in (a-b) represent significant difference in efficiency between v9h and v11h as measured by Z-score test for two proportions (*: $p<0.05$; **: $p<0.01$; ***: $p<0.001$).

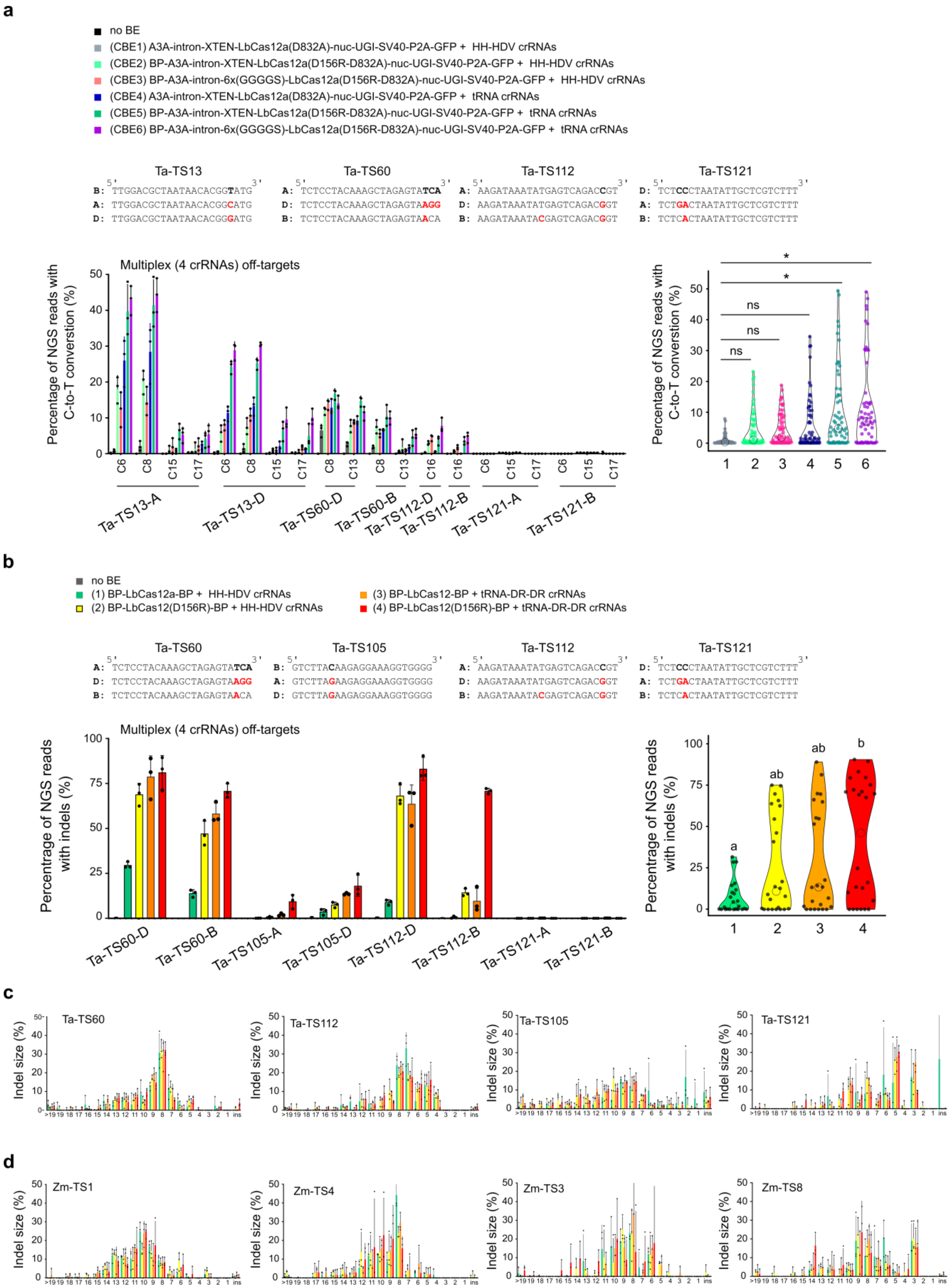
a
no BE
(CBE1) A3A-intron-XTEN-LbCas12a(D832A)-nuc-UGI-SV40-P2A-GFP + HH-HDV crRNAs
(CBE2) BP-A3A-intron-XTEN-LbCas12a(D156R-D832A)-nuc-UGI-SV40-P2A-GFP + HH-HDV crRNAs
(CBE3) BP-A3A-intron-6x(GGGGS)-LbCas12a(D156R-D832A)-nuc-UGI-SV40-P2A-GFP + HH-HDV crRNAs
(CBE4) A3A-intron-XTEN-LbCas12a(D832A)-nuc-UGI-SV40-P2A-GFP + tRNA crRNAs
(CBE5) BP-A3A-intron-XTEN-LbCas12a(D156R-D832A)-nuc-UGI-SV40-P2A-GFP + tRNA crRNAs
(CBE6) BP-A3A-intron-6x(GGGGS)-LbCas12a(D156R-D832A)-nuc-UGI-SV40-P2A-GFP + tRNA crRNAs
Ta-TS13
B: 5' TTGGACGCTAATAACACGGTATG 3'
A: TTGGACGCTAATAACACGGCATG
D: TTGGACGCTAATAACACGGGATG
Ta-TS60
A: 5' TCTCCTACAAAGCTAGAGTATCA 3'
D: TCTCCTACAAAGCTAGAGTAAGG
B: TCTCCTACAAAGCTAGAGTAACA
Ta-TS112
A: 5' AAGATAAATATGAGTCAGACCGT 3'
D: AAGATAAATATGAGTCAGACGGT
B: AAGATAAATACGAGTCAGACGGT
Ta-TS121
D: 5' TCTCCCTAATATTGCTCGTCTTT 3'
A: TCTGACTAATATTGCTCGTCTTT
B: TCTCACTAATATTGCTCGTCTTT
Multiplex (4 crRNAs) off-targets
Percentage of NGS reads with C-to-T conversion (%)
Ta-TS13-A
Ta-TS13-D
Ta-TS60-D
Ta-TS60-B
Ta-TS112-D
Ta-TS112-B
Ta-TS121-A
Ta-TS121-B
ns
*
b
no BE
(1) BP-LbCas12a-BP + HH-HDV crRNAs
(2) BP-LbCas12(D156R)-BP + HH-HDV crRNAs
(3) BP-LbCas12-BP + tRNA-DR-DR crRNAs
(4) BP-LbCas12(D156R)-BP + tRNA-DR-DR crRNAs
Ta-TS60
A: 5' TCTCCTACAAAGCTAGAGTATCA 3'
D: TCTCCTACAAAGCTAGAGTAAGG
B: TCTCCTACAAAGCTAGAGTAACA
Ta-TS105
B: 5' GTCTTACAAGAGGAAAGGTGGGG 3'
A: GTCTTAGAAGAGGAAAGGTGGGG
D: GTCTTAGAAGAGGAAAGGTGGGG
Ta-TS112
A: 5' AAGATAAATATGAGTCAGACCGT 3'
D: AAGATAAATATGAGTCAGACGGT
B: AAGATAAATACGAGTCAGACGGT
Ta-TS121
D: 5' TCTCCCTAATATTGCTCGTCTTT 3'
A: TCTGACTAATATTGCTCGTCTTT
B: TCTCACTAATATTGCTCGTCTTT
Multiplex (4 crRNAs) off-targets
Percentage of NGS reads with indels (%)
Ta-TS60-D
Ta-TS60-B
Ta-TS105-A
Ta-TS105-D
Ta-TS112-D
Ta-TS112-B
Ta-TS121-A
Ta-TS121-B
a
ab
ab
b
c
Ta-TS60
Ta-TS112
Ta-TS105
Ta-TS121
Indel size (%)
d
Zm-TS1
Zm-TS4
Zm-TS3
Zm-TS8
Indel size (%)

**Fig. S7: Cas12a-CBEs and LbCas12a efficiency at endogenous sites in protoplasts.**

**a** Base editing efficiencies of six Cas12a-CBE configurations in multiplex at four wheat off-targets located on homoeologous genes. Efficiencies are measured by the proportion of reads converted from C-to-T. DNA sequences on the top row are the on-targets, red bases indicate mismatches with the crRNA at off-target homeologs. Asterisks depicts significant difference in efficiency as calculated by Kruskal-Wallis test followed by Dunn post-hoc test (*: $p<0.05$). **b** Multiplex indel rate of LbCas12a at eight wheat off-targets located on wheat homeolog genes. DNA sequences on the top row are the on-targets, red bases indicate mismatches with the crRNA at off-target homeologs. Violin plots represent pooled indel rates at all targets for individual LbCas12a architectures. Significance is calculated by two-way ANOVA test with Tukey HSD post-hoc test at $p<0.05$. Editing rates were calculated from three independent biological replicates that are depicted as dots on barplots. **c-d** Pattern of indels generated by LbCas12a at four wheat (c) or maize (d) on-targets. Insertions and deletion from 1 to >19 base pairs are shown.

### Appendix

Raw gel pictures during amplicon isolation. Red rectangles depict PCR bands that were isolated and analyzed. Individual gels correspond to figure panels indicated on top left corner. Name of the target size and expected PCR product sizes are indicated on gel pictures.

Fig 1e

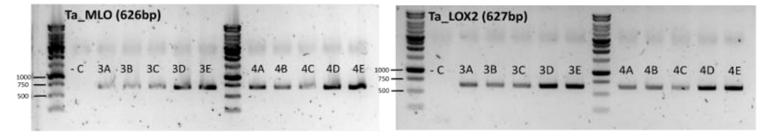


Fig 1f

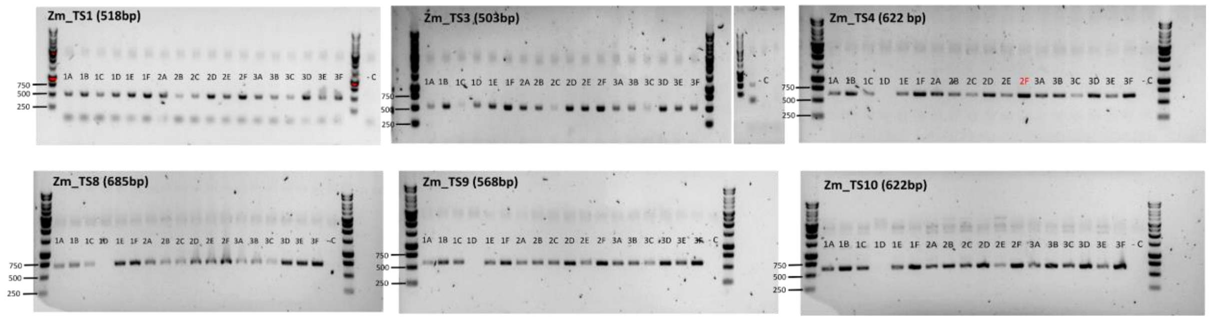


Fig 1g

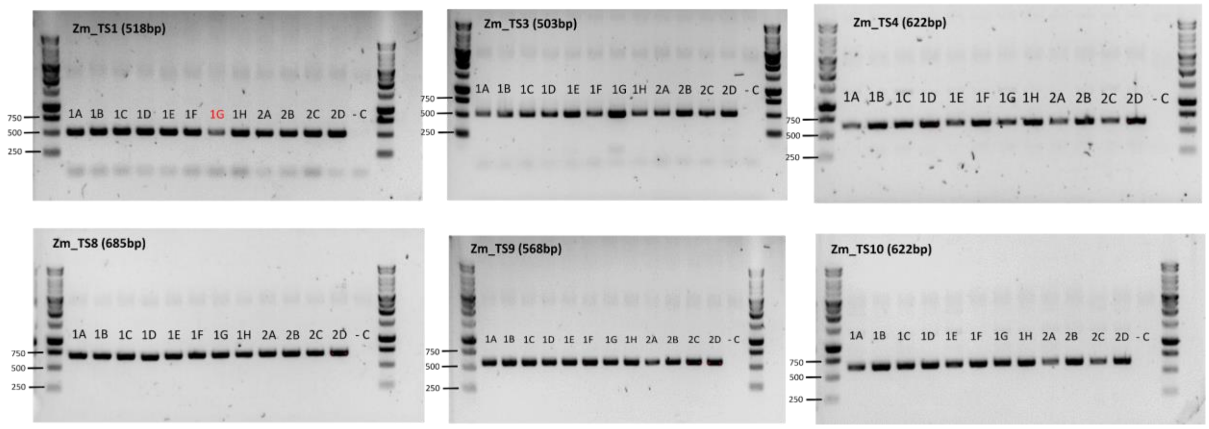

Fig 3a

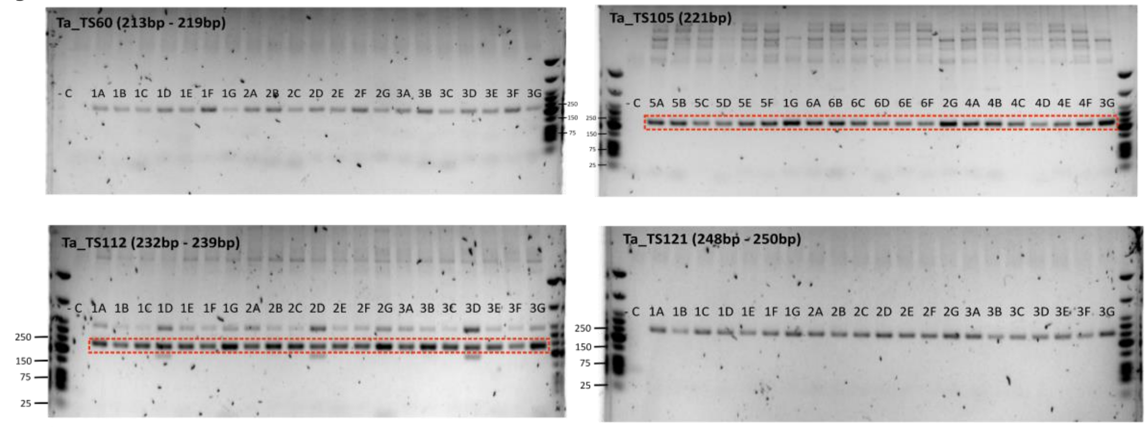


Fig 3b

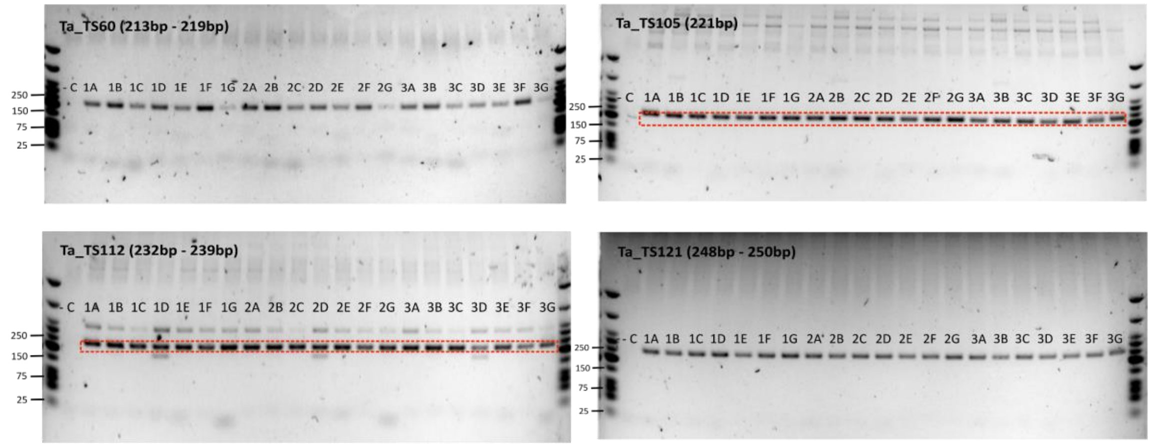


Fig 3c

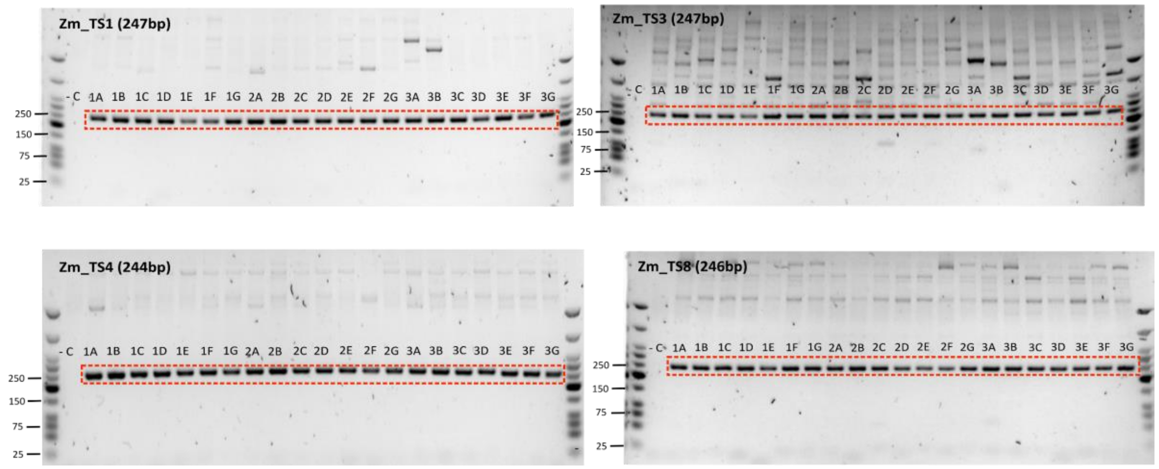

Fig 4

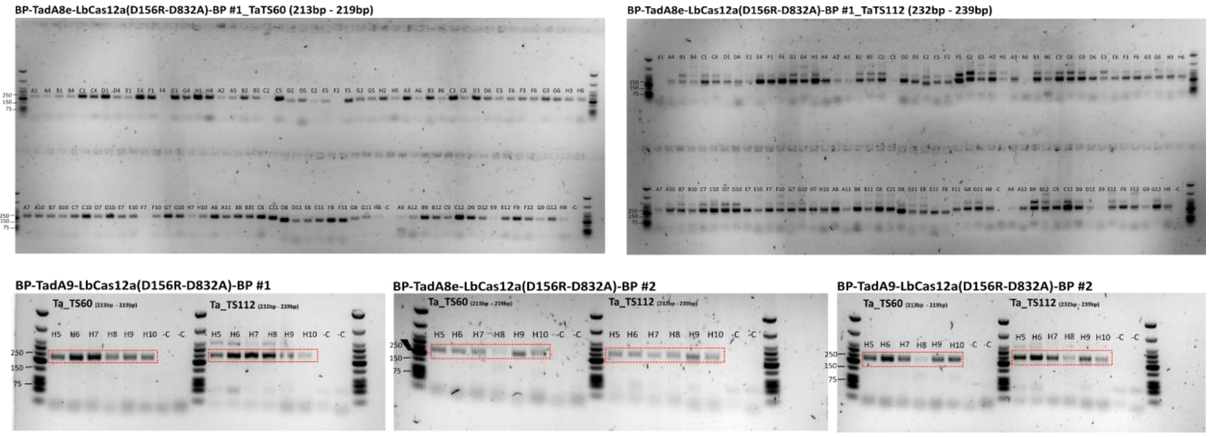
BP-TadA8e-LbCas12a(D156R-D832A)-BP #1_TaTS60 (213bp - 219bp)
BP-TadA8e-LbCas12a(D156R-D832A)-BP #1_TaTS112 (232bp - 239bp)
BP-TadA9-LbCas12a(D156R-D832A)-BP #1
BP-TadA8e-LbCas12a(D156R-D832A)-BP #2
BP-TadA9-LbCas12a(D156R-D832A)-BP #2
Ta_TS60
Ta_TS112
H5 H6 H7 H8 H9 H10 -C -C
250
150
75

Fig 5a

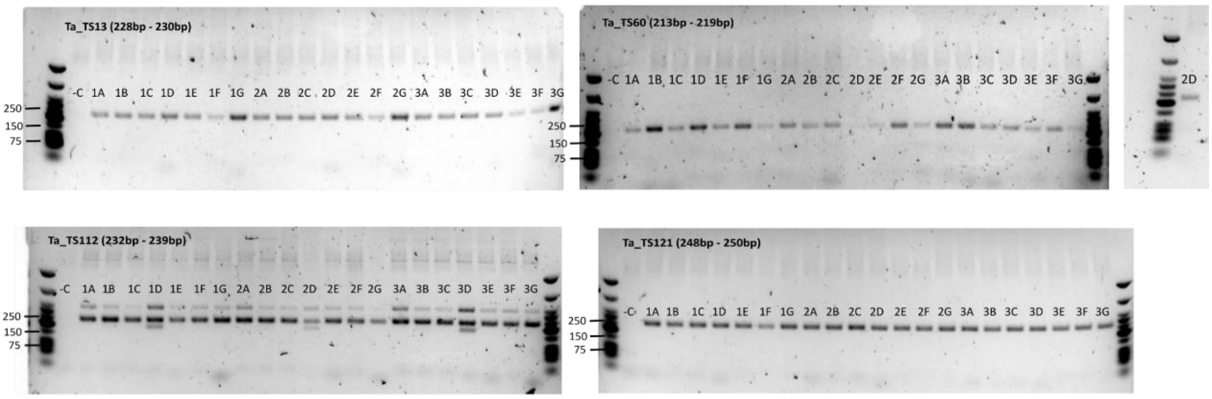


Fig 5b

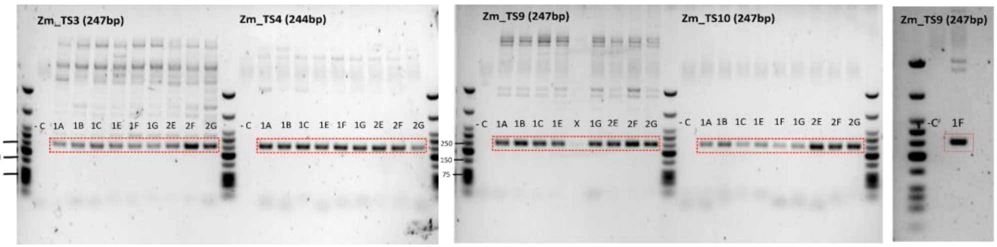


Fig 5c

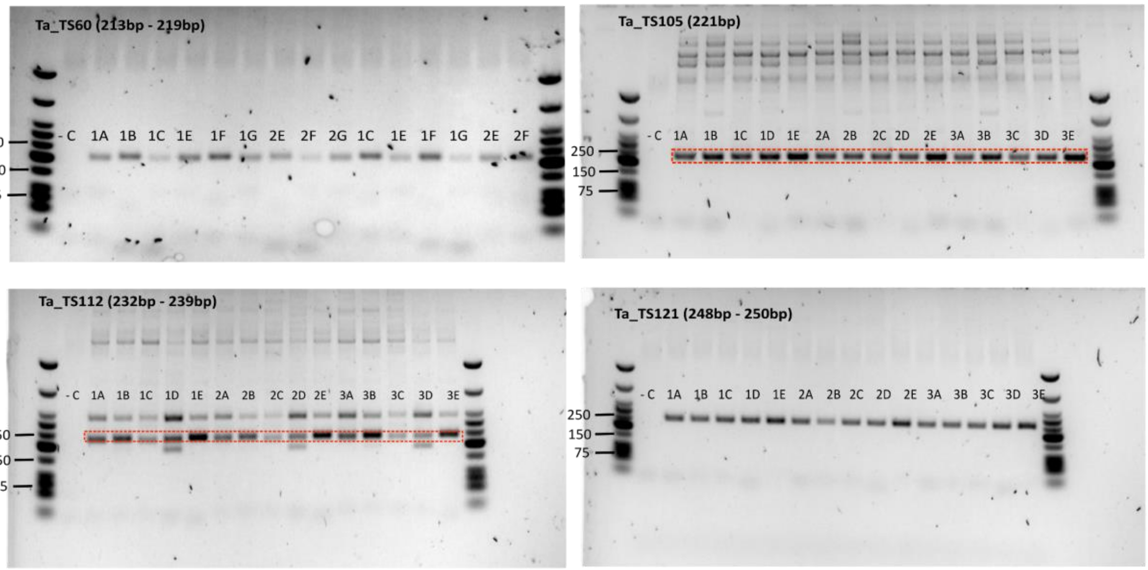


Fig 5d

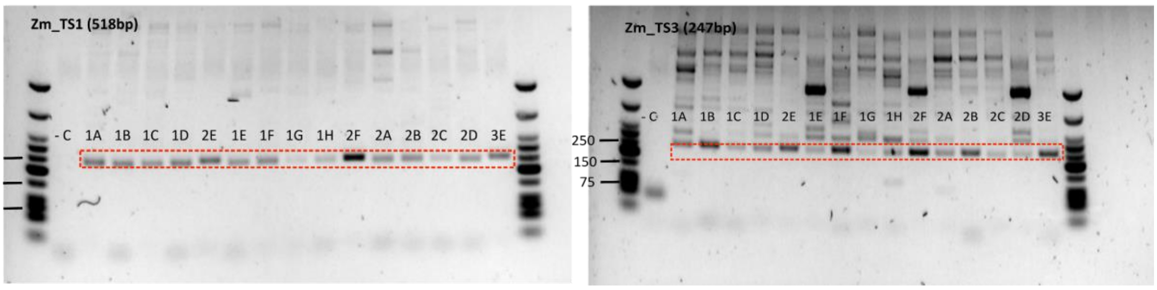

ED - Fig 3c

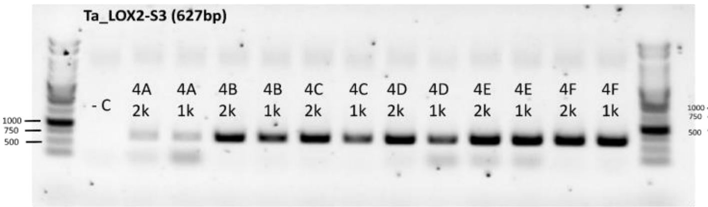


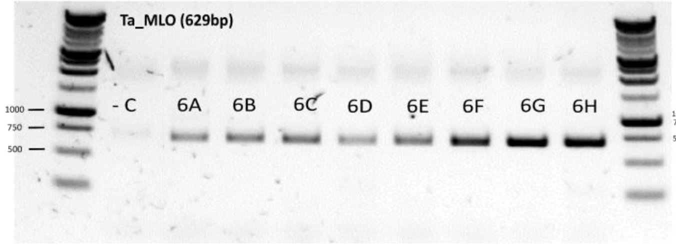


ED - Fig 3g

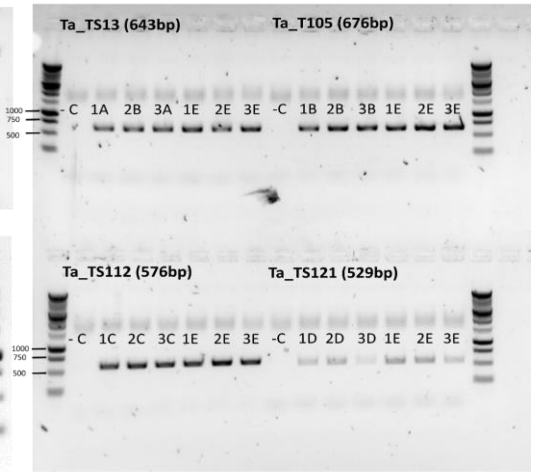


ED - Fig 5a

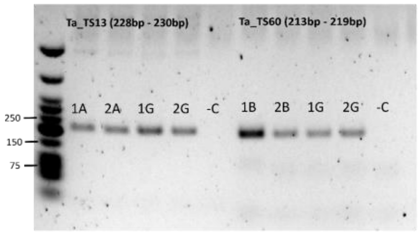


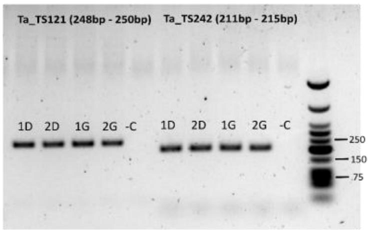


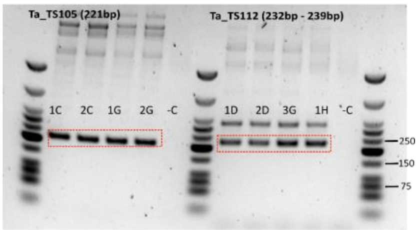


ED - Fig 5b

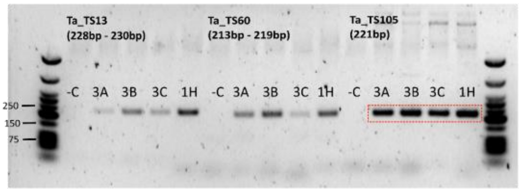


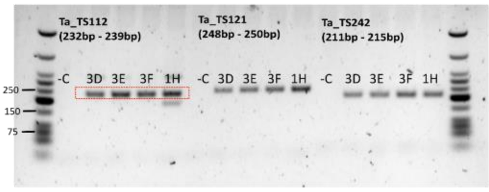


ED - Fig 5f

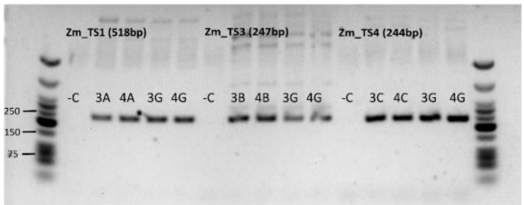


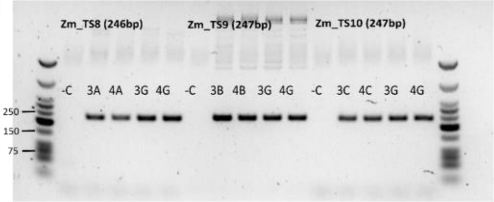


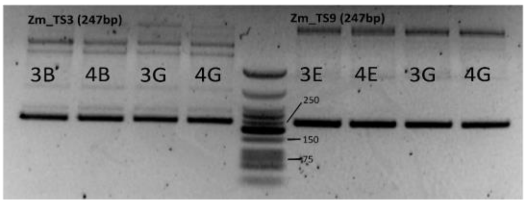
